# In-silico studies on thermodynamics of ligand binding to Fluoride riboswitch aptamer

**DOI:** 10.1101/2024.06.23.600262

**Authors:** Soumi Das

## Abstract

Riboswitch is a non-coding messenger RNA (m-RNA) whose aptamer domain binds cognate ligands and subsequently undergoes conformational changes in the expression platform leading to the regulation of gene expression. Fluoride riboswitch has immense pharmacological potential due to its presence in some human bacterial pathogens. Several experimental studies shed light on the bacterial defense mechanism of Fluoride riboswitch upon binding of F^-^ cognate ligand in the presence of Mg^2+^. However, the structural and thermodynamic basis of ligand binding with Fluoride riboswitch aptamer is not well known. This fascinates us for investigating the conformational stability of (i) the holo form of T. Petrophila fluoride riboswitch aptamer (RNA+F^-^+Mg^2+^+K^+^) with respect to (ii) the apo form of fluoride riboswitch (RNA in the absence of F^-^ +Mg^2+^+K^+^). Conformational thermodynamics results derived from molecular dynamics simulation reveal that the holo form of the Fluoride riboswitch aptamer is stabilized by ion recognition site, pseudoknot, and stem1. However, Stem2, Loop1, Loop2, and most of the unpaired bases show significant disorder and destabilization. Molecular docking study validates the thermodynamically destabilized and disordered residues from Loop1 and Stem2 of the Fluoride riboswitch aptamer to serve as putative binding sites for non-cognate ligands. The global health system in the current century faces a serious crisis to counteract bacterial infection due to the severe emergence of bacterial resistance to antibiotics. Consequently, a need for a new generation of antibiotics against resistant bacteria is critically acclaimed. Our work hopefully improves the design of new ligands and aptamers which may be helpful in nucleic acid-targeted therapeutics.

**Graphical Abstract:** 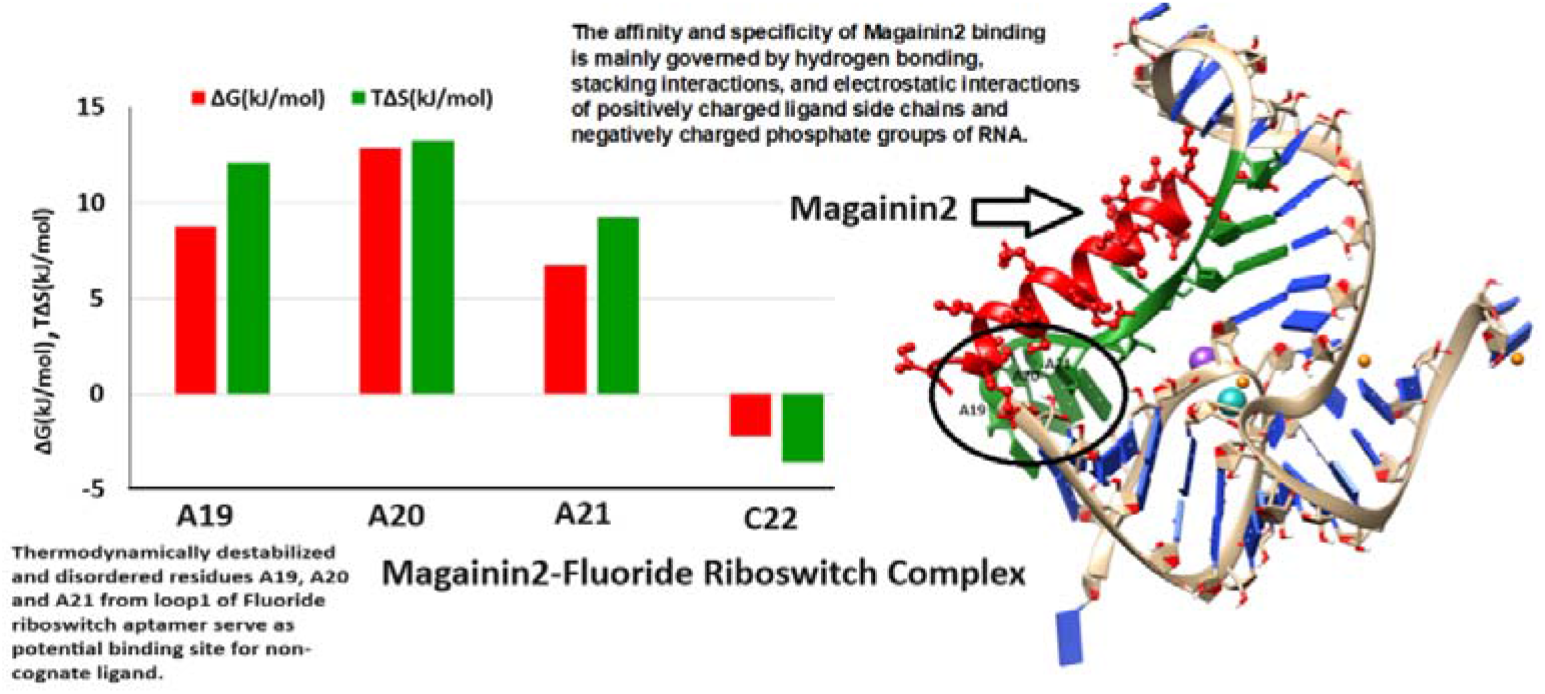

**Highlights:**

- Conformational stability of (i) the holo form of T. Petrophila fluoride riboswitch aptamer (RNA+F^-^+Mg^2+^+K^+^) with respect to (ii) the apo form of fluoride riboswitch (RNA in the absence of F^-^ +Mg^2+^+K^+^) is studied.
- Holo Fluoride riboswitch aptamer gets energetically and entropically stable at Pseudoknot, Stem1, and Ion recognition sites whereas Stem2, Loop1, Loop2, and most of the unpaired bases show significant disorder and destabilization.
- The hydrogen bond network for Pseudoknot, Stem1, and Stem2 in apo Fluoride riboswitch is significantly weak.
- Molecular Docking study confirms that the thermodynamically destabilized and disordered residues from Stem2 and Loop1 of the holo Fluoride riboswitch aptamer serve as putative binding sites for non-cognate ligands.

## 1. Introduction

Riboswitch, which resides in the 5’ untranslated region (UTR) of an mRNA, modulates gene expression in response to cognate ligand binding towards the aptamer domain. The ligand can be a metabolite or small molecule like nucleobase, amino acid, sugar, coenzyme, or metal ion. The ligand binding to the aptamer domain which induces a conformational change in the expression platform occurs through conformational selection and induced fit mechanism. The expression platform controls the regulation of gene expression via various mechanisms, like transcription, translation, and splicing **[1]**.

Fluoride riboswitch represents an example of RNA-anion interaction. The smallest and most electronegative anion F^-^ acts as a cognate ligand chelated by three Mg^2+^. The distinct antimicrobial properties of F^-^ are well known. Fluoride riboswitch plays a vital role in maintaining cytoplasmic fluoride ion concentration in the bacteria below the toxic levels. The presence of Fluoride riboswitch in several human bacterial pathogens makes it a potential drug target. It is observed that among all halide anions, only F^-^ exhibits unparalleled selectivity towards Fluoride riboswitch aptamer **[2, 3, 4]**.

**Fig. 1(a)** represents the crystal structure of the Fluoride riboswitch aptamer domain highlighting different structural regions. Fluoride riboswitch aptamer, one of the higher-order RNA architectures, is characterized by pseudoknot, stem1, and stem2 formation involving standard Watson–Crick base pairing residues connected by loop1 and loop2. **Fig. 1(b)** illustrates the coordination scheme of the cognate ligand. **Fig. 1(c)** explains how the backbone phosphate of RNA and water molecules fill up the coordination of three Mg^2+^ (Mg 1, Mg 3, and Mg 4) ions. Besides coordination scheme of the other two Mg^2+^ ions (Mg2, Mg5) and K^+^ are shown in **Fig. 1(d), Fig. 1(e), and Fig. 1(f)**. The distance cut off between the phosphate atoms of RNA and Mg^2+^ ions is set at less than or equal to 3.0 Å for the coordination scheme.

**Fig 1(a):**
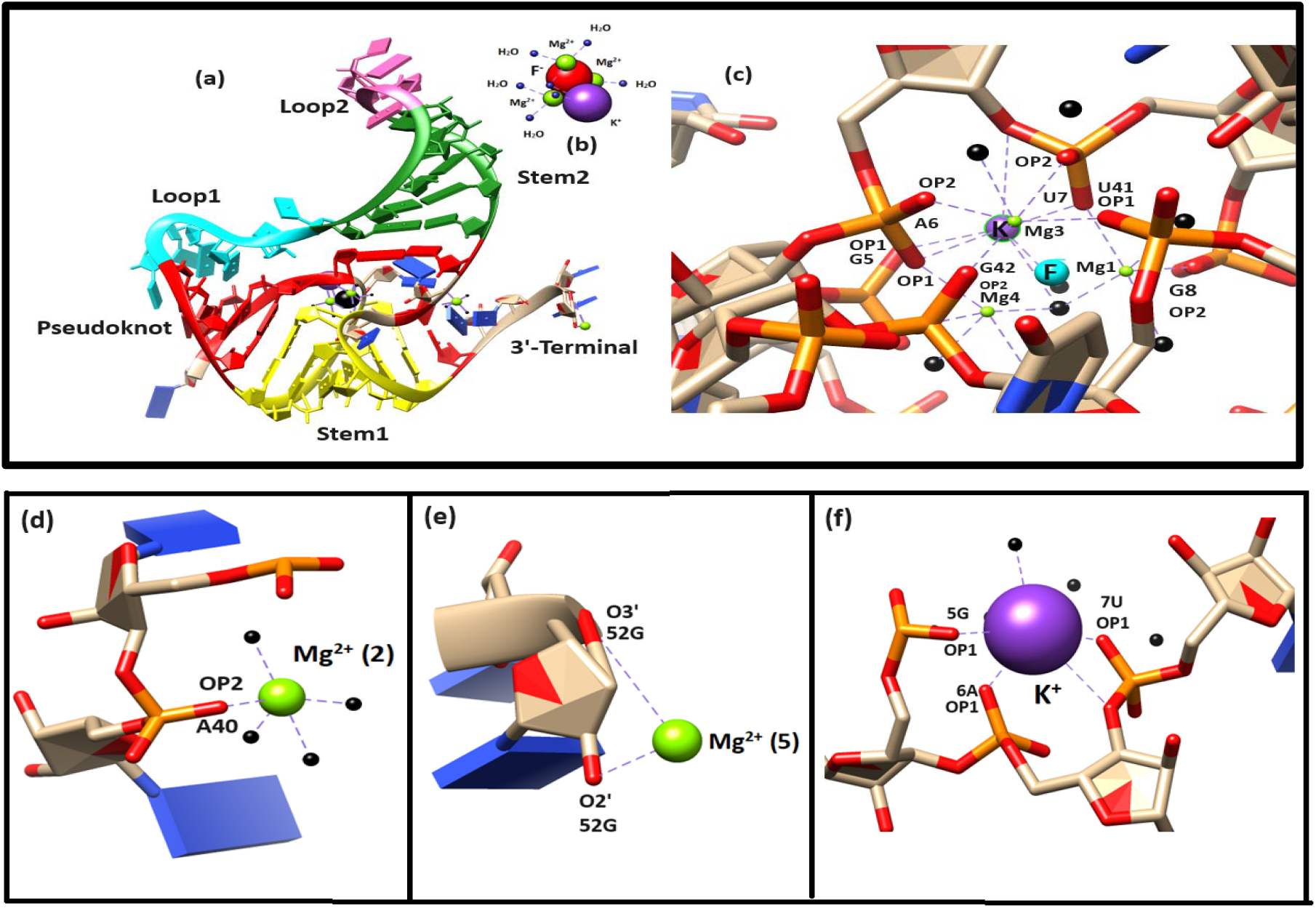
Crystal structure of the aptamer domain of Fluoride riboswitch of T. Petrophila (PDB-ID: 4ENC) with K^+^ and tri-Mg^2+^ coordination of F^-^ is shown in cartoon representation. The tertiary structural elements pseudoknot, stem1, stem2, loop1, loop2, and the 3’ terminal are shown in red, yellow, green, cyan, hot pink, and brown respectively. (b): Coordination scheme of F^-^ (red), Mg^2+^ (green), and K^+^ (violet). The blue sphere indicates coordinated water. (c): Coordinating residues of Mg^2+^of 1, 3, and 4 (d): Coordinating residues of Mg^2+^ of 2 (e): Coordinating residues of Mg^2+^of 5 and (f): Coordinating residues of K^+^. The black sphere indicates coordinated water.

SHAPE-seq, CESTNMR, and sm-FRET studies on the Bacillus cerus crcB Fluoride riboswitch indicate the stability of Fluoride riboswitch aptamer in different ion states **[5, 6]**. Although the experimental structure of the Fluoride riboswitch aptamer is now available, details structural and thermodynamic basis of the interaction of the cognate ligand binding to Fluoride riboswitch aptamer are missing in the literature. Binding free energy and binding entropy measurement using isothermal titration calorimetry can not yield region-wise information on conformational change. To fill this void, we examine the conformational stability and order of holo Fluoride riboswitch aptamer with respect to the apo form using conformational thermodynamics technique derived from equilibrium trajectories of atomistic molecular dynamics simulation **[7]**. It is a necessary step for drug design to derive the thermodynamic basis of biomolecular interaction elaborately for interpreting key residues involved in binding event. Thermodynamics of region-wise conformational change of Fluoride riboswitch aptamer upon F^-^ cognate ligand binding is extracted from the histograms of microscopic conformational variables. The histogram-based method (HBM) for conformational thermodynamics estimation which was initially developed for the proteins **[8, 9, 10, 11, 12]** further extended to the protein-DNA complex systems as well **[13, 14]**. Here we further extend the conformational thermodynamics technique to the RNA system using the microscopic conformational variables of RNA, described in SI are (i) inter-bp step parameters: tilt (τ), roll (ρ), twist (ω), shift (D_x_), slide (D_y_), and rise (D_z_). (ii) intra-bp step parameters: buckle (κ), open (σ), propeller (π), stagger (S_x_), shear (S_y_), stretch (S_z_). (iii) sugar-phosphate backbone torsion angles: α, β, γ, δ, ε, ζ and sugar-base backbone torsion angle χ. (iv) sugar pucker: (v_0_, v_1_, v_2_, v_3_, v_4_) and (v) pseudo torsion angle (□ and θ). Conformational stability and order of RNA-ligand complex with respect to free RNA are quantified by negative changes in conformational free energy and entropy while positive changes demonstrate conformational destabilization and disorder respectively. The inclusion of the expression platform of Fluoride riboswitch in molecular dynamics simulation is difficult due to the scarcity of structural information because the expression platform cannot be crystallized due to high fluctuations. Removal of the ligands (F^-^, Mg^2+^, and K^+^) leads to significant distortions of the phosphate backbone near the ligand binding pocket interpreted from the fluctuations of microscopic conformational variables. Our conformational thermodynamics data show that the ion recognition site, pseudoknot and stem1 are significantly stabilized and ordered whereas loop1, loop2, and stem2 get completely destabilized and disordered among all other regions of fluoride riboswitch aptamer. The residues of the holo form having instability in conformational free energy and disorder in entropy are supposed to be involved in additional binding events to attain thermodynamic stability and order by reducing the free energy and entropy. The thermodynamically destabilized and disordered residues from stem2 and loop2 of Fluoride riboswitch aptamer are suggested to act as binding sites for non-cognate ligands. This hypothesis is examined for two known antibiotics Gramicidin D and Magainin2 binding to Fluoride riboswitch via docking. Conformational thermodynamics data and docking results ensure that destabilized and disordered residues of loop1 and stem2 of holo Fluoride riboswitch act as receptor interfaces. Our work can be generalized to underpin conformational aspects, binding mode, and energetics of the non-cognate ligand binding to other riboswitches. Key findings from this study would hopefully benefit nucleic acid-targeted therapeutic research areas.

## 2. Methods

### 2.1 System preparations

The holo T. petrophila Fluoride riboswitch aptamer coordinates are taken from PDB-ID 4ENC. Two different cases of the aptamers are modeled:

| System | Duration of Molecular dynamics simulation | Force field | RMSD (mean and SD) |
| --- | --- | --- | --- |
| Holo form of fluoride riboswitch<br>(PDB-ID: 4ENC)<br>(RNA + F<sup>-</sup> + Mg<sup>2+</sup> + K<sup>+</sup>) | 1 μs | $\chi_{OL3}$ RNA Force field | 3.98 Å(0.05) |
| Apo form of fluoride riboswitch<br>(Without F<sup>-</sup> + Mg<sup>2+</sup> + K<sup>+</sup>) | 1 μs | $\chi_{OL3}$ RNA Force field | 2.2 (0.07) |

### 2.2 Simulation details

All MD simulations are performed using the GROMACS simulation program **[15]** with χOL3 force-field parameters for the RNA **[16]**. For explicit solvent calculation, the RNA systems with and without the ligand are solvated in a cubic box with 15 Å thick layers of water molecules and neutralized with the addition of the required number of sodium [Na^+^] and chloride [Cl^-^] ions. The periodic boundary conditions are imposed in all directions. Subsequently, the systems are energy minimized using the steepest descent algorithm **[17]**. Particle Mesh Ewald summation (PME) is used for long-ranged electrostatic interaction with 1 Å grid spacing and 10^-6^ convergence criterion. The Lennard-Jones and the short-range electrostatic interactions are truncated at 10 Å. The all-atom molecular dynamics (MD) simulation is carried out at 300K temperature and 1 atmospheric pressure in an isothermal-isobaric (NPT) ensemble starting from the energy minimized structure. Berendsen thermostat **[18]** is used to keep up a constant temperature and the pressure is controlled by the Parrinello-Rahman barostat **[19]**. The LINCS constraints are applied to all bonds involving hydrogen atoms. Integration time step of 1 fs is used. We perform MD simulation for holo and apo Fluoride riboswitch aptamer with 1 µs duration. The trajectories are visualized employing VMD program and images are captured with the help of CHIMERA **[20]** and PYMOL **[21]**.

### 2.3 Structure analysis

The equilibrations for different systems are judged from the root mean square deviations (RMSD). The backbone phosphorous (P) atom based root mean square fluctuation (RMSF) per residue is computed to further study the flexibility of each nucleotide and compare the difference in dynamics between unliganded and liganded simulations. Different inter-base and intra-base pair parameters, torsion angle, pseudo-torsion angle, and stacking overlap are calculated using NUPARM software **[22]**. Base pairing information is detected by BPFIND **[23]**.

### 2.4 Identification of interactions

The interactions are characterized by distance and angle criteria. The hydrogen bonds are taken when the distance between the donor (D) and acceptor (A) is less or equal to 3.5 Å and the angle (D-H-A) cut-off is 160^0^. GROMACS Hydrogen bond analysis module is used for hydrogen bond network calculation. Electrostatic interaction is considered when the distance between oppositely charged atoms is less than 5.6 Å.

### 2.5 Conformational thermodynamics

The detailed description of the histogram-based method (HBM) for calculating the conformational thermodynamics is reported **[8, 9, 10, 11, 12, 13, 14]**. The histograms refer to the probability of finding the system in a given conformation. The histograms can be interpreted with the help of the Boltzmann factors corresponding to effective free energy and entropy is estimated by Gibbs Formula. The conformational thermodynamics and the histograms are interconnected in this way. The computations are done using the code in https://github.com/snbsoftmatter/confthermo. The histograms of microscopic conformational variables are computed using equilibrated trajectory, namely, 600 to 1000 ns. The normalized probability distribution of any microscopic conformational variable θ in the free state and the complex state are given by H^free^ (θ) and H^complex^ (θ), respectively. The change in free energy of any microscopic conformational variable θ of the bound state as compared to the free state is defined as,

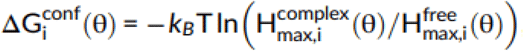

where “max” represents the peak value of the histogram and i represents the RNA residue.

The change in conformational entropy of a given microscopic conformational variable θ of the bound state as compared to the free state is evaluated as,

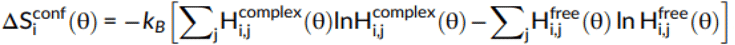

where the sum is taken over all histogram bins j and i represents the RNA residue.

### 2.6 Docking studies

The average structure of holo Fluoride riboswitch calculated from equilibrated MD trajectory is used as the receptor for docking procedure while peptide drugs Gramicidin D and Magainin 2 as ligands. The coordinates of Gramicidin D is extracted from PDB-ID: 1BDW. The coordinates of Magainin 2 is segregated from PDB-ID: 2MAG. At first molecular docking webserver HDOCK **[24]** specialized in protein-protein and protein-DNA/RNA interactions is used. Next HADDOCK **[25]** is used for docking to compare the docking score between two structures for two cases. The docking protocol encompasses three stages namely rigid body energy minimization, semi-flexible simulated annealing in torsion angle space, and explicit solvent refinement. We bias the docking procedure by considering destabilized and disordered residues of loop1 and stem2 of Fluoride riboswitch aptamer as active residues for binding. The resulting docked structures are sorted by minimum energy criteria and RMSD clustering. The central structure of the cluster with the maximum z score is selected as the best docked structure. Next, the interface of the docked complex is analyzed based on the distance cut off by 5 Å between residues from the binding partners. We further minimize the docked complexes in the explicit solvent imposing the steepest descent algorithm, until the maximum force on any atom is less than 100 kJ mol^-1^ nm^-1^ with the Amber protein force field ff14SB for the peptides and χ_OL3_ force-field parameter for the RNA.

- **Abbreviations of non-canonical base pair**

The **W: WT** base pair involves Watson-Crick edges of both the bases but in trans orientation. This is traditionally called the reverse Watson-Crick base pair.

The **H: WT** base pair involves the Hoogsteen edge of the first base pairing with the Watson-Crick edges of the second base in trans orientation.

The **s: sC** base pair involves the sugar edge of the first base pairing with the sugar edge of the second base in cis orientation. The base pair has one week non-polar C-H…N hydrogen bond.

The **s: sT** base pair involves the sugar edge of the first base pairing with the sugar edge of the second base in trans orientation. The base pair has one week non-polar C-H…N hydrogen bond.

The **s: hT** base pair involves the sugar edge of the first base pairing with the Hoogsteen edge of the second base in trans orientation. The base pair has one week non-polar C-H…N hydrogen bond.

## 3. Results

The equilibrated snapshots of the holo and apo fluoride riboswitch at 1000 ns are shown in **Fig. 2(a) and (b)** respectively. RMSDs of both the holo and apo structure **(Fig. S1 (a))** show variations in RMSD values in an acceptable range; the average RMSD for ligand-free apo structure is 4.1 Å whereas for the ligand-bound holo-structure that is 3.98Å. The equilibrated part of the trajectory, (600ns-10000ns), is considered for further analysis.

**Fig. 2.**
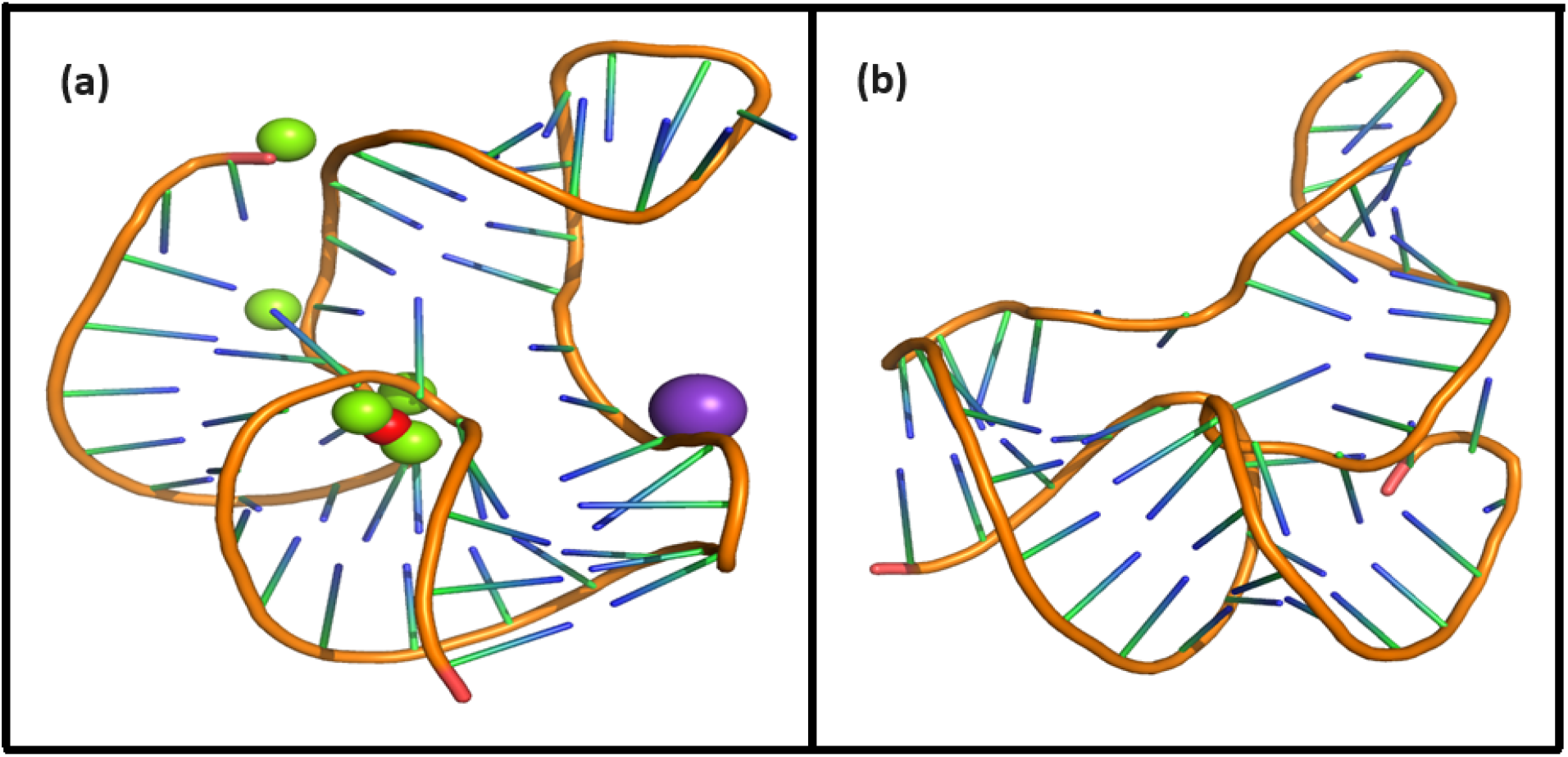
(a) and (b): Snapshot at 1000 ns of holo and apo aptamer domain of fluoride riboswitch respectively.

### 3.1 Global and local dynamics of the aptamer domain and Interaction

The stem1, stem2, pseudoknot, and ligand recognition sites show lower fluctuations for both ligand-free and ligand-bound simulations as reflected from root mean square fluctuations (RMSF) **(Fig. S1 (b))**. Fluctuations in loop1 (residues 18, 19, 20, 21, 22), loop2 (29, 30, 31, 32), and terminal regions (49, 50, 51, 52) are in general higher than in other regions of the aptamer. The free motion of the terminal regions of the aptamer results in relatively higher fluctuations. The fluctuations become more pronounced in apo form with a sharp peak indicating highly dynamic and flexible loop region. Fluctuations are the least in the ion binding sites (residues 5, 6, 7, 40, 41, 42). On a local scale, local conformational rearrangements in the loop region are manifested by higher RMSF values.

**Table S1 (b)** compares the probabilities of hydrogen bond interactions in the pseudoknot, stem1, and stem2 regions for the holo and apo system. Donor-acceptor distance and the donor-hydrogen-acceptor angle cut-off are set as 3.4 Å and 120^°^ respectively. Hydrogen bond probabilities P_HB_> 80% represent Strong (s) hydrogen bonds while 40% <P_HB_ <80% correspond to Weak (w) hydrogen bonds. The number of Hydrogen bond is higher in holo form compared to apo form. We find that the presence of Mg^2+^ and F^-^ significantly strengthen almost all Watson-Crick base pair hydrogen bonding interactions in the pseudoknot and stem1 of the holo form. F^-^ ligand binding to the aptamer domain imposes stabilization of the pseudoknot and inhibits the formation of terminator hairpin leading to the transcription of the downstream gene (ON state). On the other hand, weak hydrogen bonds are observed in the holo and apo form of stem2. Watson-Crick base pairing in pseudoknot and stem1 are frequently ruptured in the apo form due to the formation of a transcription terminator (OFF state) involving a stable stem-loop conformation. There occurs a conformational change of Fluoride riboswitch aptamer in the absence of cognate ligand F^-^. Hydrogen bond analysis thus confirms that the aptamer domain and expression platform of the T. Petrophila Fluoride riboswitch interconverts between two conformations.

The coordination sphere of the ligand binding site is analyzed by the geometric configuration in terms of bond length in the holo system (**Table S1(c), Fig. S2, S3, S4, and S5)**. It is observed from distance calculation over the MD trajectory that the interaction between Mg1-F and Mg4-F is higher compared to Mg2-F, Mg3-F and Mg5-F. Average smaller distance (**2Å**) of Mg1-OP1 (7U), Mg1-OP2 (8G), Mg3-OP2 (6A), Mg3-OP2 (7U), Mg3-OP1 (41U), Mg3-OP2 (42G), Mg4-OP1 (6A) and Mg4-OP1 (42G), indicate strong electrostatic interaction. The distance between Mg^2+^ (1, 2, 3, 4, and 5) and F^-^ (**Fig. S2 (a), (b), (c), (d) and (e))**, as well as Mg^2+^ and the backbone of Fluoride riboswitch (**Fig. S3, S4, and S5)**, remain nearly constant throughout the simulation time indicating stable interaction. Mg^2+^-backbone distances suggest that Mg^2+^ being an essential constituent of the fluoride riboswitch forms stable metal coordination site. On the other hand **Table S1(c)** shows that the interaction between K^+^ and the backbone of Fluoride riboswitch is highly unstable and dynamic. There is no direct interaction between F^-^ and backbone phosphates of fluoride riboswitch aptamer. The F^-^ ion is coordinated to three Mg^2+^ ions (Mg1, Mg3, and Mg4), which in turn are chelated to G5pA6pU7pG8 and A40pU41pG42 backbone phosphates and water molecules **(Fig. 1(c))**.

### 3.2 Structural variability

Base pair parameters and base pair step parameters deliver a great deal of information to assess the overall geometry of the double-helical region of fluoride riboswitch such as pseudoknot, stem1, and stem2 **(Tables S2 (a) and (b), and S3 (a) and (b))**. Roll values are mostly observed as large positive (∼10^°^). In most cases, twist values are around 30^0^–40^0^. Slide values are around -1.5 Å in A-form RNA. Rise values are around 3.4 Å in RNA. Base pair parameters and base pair step parameters suggest that most of the base pairs in the pseudoknot and stem regions of holo and apo form of Fluoride riboswitch aptamer predominantly adopt A-form RNA structures with proper stacking between canonical base pairs. Standard deviation values are within acceptable ranges for most of the base pairs, indicating reasonable conformational stabilities. Average roll and twist angles are rather unusual suggesting deformation of some of the steps involving canonical base 5 G:C14 W:W C (pseudoknot), 28 U:A 33 W:W C (stem2) as well as non-canonical base-pairs 21 A:G 3 s:s T (loop1), 38 U:A 6W:W T (pseudoknot) in holo form. On the other hand 20 A:C 17s:s C (loop1), 28 U:A 33 W:W C (stem2), 38 U:A 6 W:W T (pseudoknot) of apo form exhibit significant conformational alteration. Slide value ranges from small positive values to as large as -3 Å for the steps containing non-canonical base-pairs. Buckle, propeller, and stagger values indicate high stability with strong, stable, and planar base pairing of most of the base-pairs in pseudoknot and stem1 of holo and apo form against deformation or rupture of H-bonds. Large open angle is indicative of disruption of the H-bond in loop1 of 20 A:Cs:s C 17, 21 A:Gs:s T 3, and ion recognition site of 41 U:Cs:h T44 for both holo and apo.

**Tables S2 (c) and S3 (c)** show the backbone torsion angles: χ; pseudo-rotation phase angle (P) and pseudo-torsion angles: □ and □. The glycosidic torsion angle χ adopts an anti-conformation (+90^0^ to +180^0^; –90^0^ to –180^0^) whereas sugar pucker prefers C3’-endo conformation for most of the residues within [0^0^, 36^0^] range for fluoride riboswitch aptamer in all two systems. Our data show that A6 (pseudoknot), A19 (loop1), G24 (stem2), G42 (stem1), G52 (terminal residue) of holo and G24 (stem2), G42(stem1), G52 (terminal residue) of apo adopt syn conformation (-90^0^ to+90^0^). η and θ values are clustered around −170^0^ (-170^0^+360^0^=190^0^) and −140^0^ (-140^0^+360^0^=220^0^) respectively indicating a regular helical-like conformation of pseudoknot, stem1, and stem2. Unusual average □ and □ values are observed in loop and terminal regions for both holo and apo systems with often very high standard deviations. They exhibit C2’-endo pseudorotamer to accommodate the conformational strain. The pseudo-rotation phase angles for all the bases of loop1 and the terminal region show a bimodal or multimodal distribution, indicating higher structural variability.

**Table S4** represents the stacking overlap values between two Watson-Crick base pairs in the helical region of pseudoknot, stem1, and stem2 which are found to be around 40-55 Å^2^ indicating significant stacking. Medium amount of stacking is observed between isolated bases in loop1 and loop2. However, non-canonical base pair 20 A: C 17 (s:s C) and 21 A: G 3 (s:s T) of loop1 do not stack well.

### 3.3 Conformational thermodynamics

We show histograms (**Fig. 3**) for a few conformational variables in different RNA conformations. The most probable value of the corresponding microscopic conformational variable is denoted by the peak of the histogram. **H(ζ**_**G5**_) and **H(ζ**_**A6**_), the histogram of **ζ** for base G5 **(Fig. 3(a))** and α for base A6 **(Fig. 3(b))** of pseudoknot of holo form show that the height of the single peaked distribution in holo form is higher than that of the apo form indicating flexibility of Ion recognition site G5 and A6 gets significantly reduced in holo form. In **Fig. 3(c), H(**β_**A20**_) of loop1 shows a single peak in apo form whereas the single peak changes to multi peaks in holo form. In contrast double peak of apo form in **H(δ**_**C22**_) of loop1 reverts back to a single peak in holo form reflected from **Fig. 3(d)**. Again we see **H(**v**3**_**G36**_) of stem2 (**Fig. 3(e)**) shows single peak in both holo and apo form, but the height of the peak in holo form is higher than that of apo form. Single broad peaked distribution in ion recognition sites **H(**□_**U41**_) and **H**(η_**G42**_) in the both of the apo form suggest significant randomness (**Fig. 3(f) and 3(g)**). On the other hand (**Fig. 3(f) and 3(g)**) sharp single peaked distribution in the holo form indicate reduced conformational fluctuation in ion recognition sites. Distributions of □ for C50 (**Fig. 3(h)**) of terminal residue **H(**□ _**C50**_) are bimodal in both states indicating large flexibility. The sharper peaks of the holo form in the majority of cases confirm that ligand binding reduces the flexibility of the Fluoride riboswitch. Severe conformational fluctuations are illustrated by the large width of the peaks. Bimodal distribution sheds light on increased fluctuations between two isomeric conformations whereas sharp single peak distribution suggests conformational stability in single isomeric conformation with reduced randomness. The changes in free energy and conformational entropy are computed considering the histograms of all conformational microscopic degrees of freedom for the holo form of Fluoride riboswitch aptamer with respect to the apo form of Fluoride riboswitch aptamer. Next we discuss region wise total changes in conformational thermodynamics (kJ/mol) of Fluoride riboswitch.

**Fig. 3.**
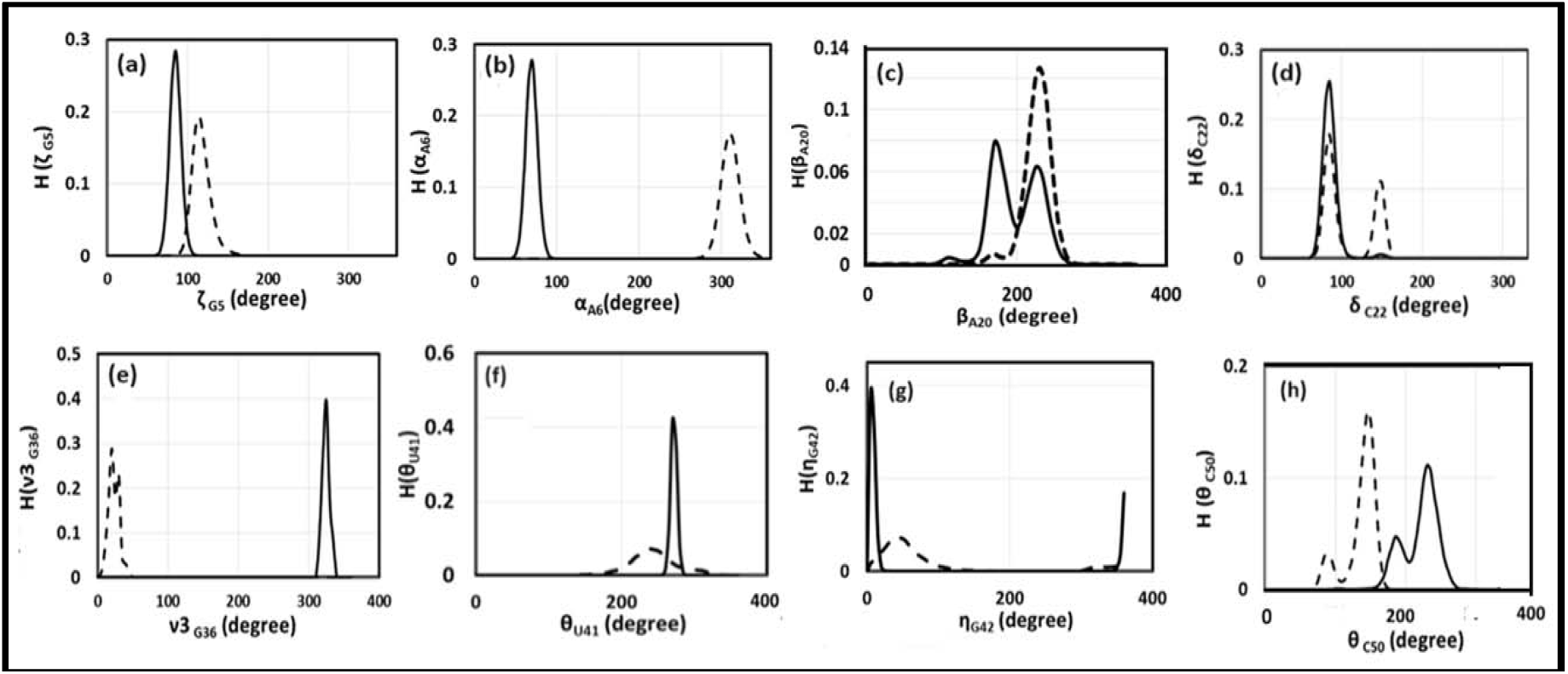
Histograms of torsion angles: (a) ζ of base G5, (b) α of base A6, (c) β of base A20, (d) δ of base C22, (e) v3 of base G36 and pseudo torsion angles: (f) □ of base U41, (g) η of base G42, (h) □ of base C50 for holo and apo systems in equilibrated trajectories. The solid line indicates the holo system whereas the broken line symbolizes the apo system.

#### Pseudoknot

We first consider the conformational thermodynamic changes in free energy ΔG and entropy TΔS caused by inter-bp step parameters (**Fig S7 (a) and (b))**. Tilt, roll, shift, and rise corresponding to the pseudoknot region do not depend significantly on the base pair steps. On the other hand, twist has maximum stabilization at step G5:C14 and destabilization at A6:U38. We observe that TΔS for all the inter base-pair step parameters except slide is not sensitive to base pair steps. The step A6:U38 shows the maximum order by slide. The total changes in conformational free energy and entropy for each bp of inter-bp parameters exhibit the highest stabilization and order at A40:U48. Next, we consider the intra-bp step parameters **(Fig S7 (c) and (d))**. Open, propeller, shear, stagger, and stretch corresponding to the pseudoknot region are not sensitive to bases. The buckle value at A6:U38 exhibits maximum stabilization and order. Finally, total changes in conformational free energy and entropy are computed for each bp of intra-bp parameters. It shows that significant conformational destabilization and disorder at G2:C17, G3:C16, and A40:U48 while maximum stabilization and order at A6:U38.

Torsion angle: α, β, γ, **δ**, ε, **ζ** and χ do not contribute significantly to bases of pseudoknot **(Fig S7 (e) and (f))** except at A6:U38. A6 and U38 show maximum order and stabilization corresponding to the evaluation of the total changes in conformational free energy and entropy over all degrees of freedom of the torsion angle. Next, we shed light on the changes in conformational thermodynamics in terms of pseudo torsion angles **(Fig S7 (g) and (h))**. ΔG of η and θ at A6 corresponding to (i) holo form with respect to apo form show maximum stabilization. ΔG and TΔS of θ at C18 contribute maximum destabilization and disorder for holo form with respect to apo. ΔG of η does not contribute significantly to bases from G15 to C18. We observe that A6 and U38 get highly stabilized and ordered based on the computation of total changes in conformational free energy and entropy in terms of pseudo-torsion angle.

Let us now shed light on the total changes in conformational thermodynamics due to sugar-pucker at pseudoknot **(Fig S7 (i) and (j))**. □0, □1, □2, □3 and □4 do not contribute significantly towards G2, G3, C16 and C17.□0 exhibits stabilization and order at C4, G5, C14and G15.□1 shows the significant stabilization and order at bases G5 & C14 whereas □2 exhibits the highest stabilization and order at A6 and U38. On the other hand, we find that □0 and □4has maximum destabilization at A6, and U38.□3 shows maximum stabilization at base C4 and G15. We evaluate the total changes in conformational free energy and entropy over the sugar torsion angle (□0, □1, □2, □3, and □4) which shows the significant stabilization and order at the bases G5, A6, C14, U38, A40, U48 while the most destabilization and disorder is observed at the bases G2, G3, C16 and C17.

#### Stem1

At first, we shed light on the changes in conformational thermodynamics in terms of inter-bp step parameters (**Fig S8 (a) and (b))**. Tilt, roll, twist, and shift corresponding to stem1 shows maximum stabilization at the step C12:G43 whereas the steps G8:C47, and A9:U46 are energetically destabilized by roll, twist, and slide. Let us now consider the changes in entropy. TΔS of tilt, roll, slide, and shift are not sensitive to base steps A9:U46 and G10:C45. On the other hand, rise has most disorder at base step 11G:C44 and C12:G43. Now we evaluate the total changes in conformational free energy and entropy for each bp due to inter-bp step parameters. C12:G43 exhibits maximum stabilization while maximum destabilization occurs at G10:C45. Total changes in conformation entropy show significant conformational order at G8:C47, C12:G43, and C13:G42. Next, we discuss the changes in conformational thermodynamics due to intra-bp parameters (**Fig S8 (c) and (d))**. The conformational free energy shows that open contributes the most to stabilizing the base step C12:G43 while buckle causes the maximum destabilization at step 11G:C44. Stretch is not sensitive to base steps. ΔG and TΔS of buckle, propeller and shear contribute the most to stabilizing A9:U46. Evaluation of the total changes in conformational free energy and entropy over the intra-bp parameters exhibits maximum order and stabilization at A9:U46.

ΔG and TΔS of torsion angle do not show sensitivity towards the bases for holo system **(Fig S8 (e) and (f))**. Pseudo-torsion angle (η and θ) of ΔG and TΔS do not show sensitivity towards the bases for holo system except G42 **(Fig S8 (g) and (h))**. Next we consider the changes in conformational free energy for sugar-puckers **(Fig S8 (i) and (j))**. v1, v2 and v4 exhibit significant stabilization at C12 and G43. TΔS are not sensitive to the bases. Computation of the total changes in conformational free energy and entropy indicates the highest disorder and destabilization at A9 and U46 while maximum order and stabilization is observed at C12 and G43.

#### Stem2

ΔG for inter-base pair step parameters corresponding to stem2 **(Fig S9 (a) and (b))** are not sensitive to the base pair steps. On the other hand, twist contributes to maximum conformational disorder at the step U28:A33. Next we consider the intra-bp parameters (**Fig S9 (c) and (d))**. The changes in conformational free energy and entropy for the holo system with respect to the apo form do not depend significantly to the bps. The changes in conformational thermodynamics for sugar-phosphate and sugar-base torsion angles are not sensitive to the bases **(Fig S9 (e) and (f))**. The total changes in conformational free energy and entropy indicate that G24, C37, C25, and G36 bases have significant stability and order. ΔG of η has maximum destabilization at U28 and A33. We observe that ΔG of η and θ are not sensitive to the bases from G34 to C37. ΔG and TΔS of θ show destabilization and disorder at C27, U28, and A33 with a maximum value at C27 **(Fig S9 (g) and (h))**. Let us now dicuss on the changes in conformational thermodynamics due to sugar-puckers **(Fig S9 (i) and (j))**. The changes in conformational free energy due to □0 indicate stability of the bases C25, G36, C26, G35, C27, G34. □1, □2, □3 and □4 exhibits destabilization towards all bases belong to stem2. Computation of the total changes in conformational free energy exhibits maximum stabilization at bases C25 and G36 and maximum destabilization at bases C27 and G34. We note that TΔS due sugar-pucker are not sensitive to bases. Total changes in conformational entropy shows maximum order at base C25 and G36 while the maximum disorder is observed at C27 and G34.

#### Loop1

We see that ΔG and TΔS of sugar-phosphate backbone torsion angles along with sugar-base torsion angle at all bases corresponding to loop1 are not sensitive to bases **(Fig S10 (a) and (b))**. Calculation of the total changes in conformational free energy and entropy exhibit the most conformational destabilization and disorder at the base A20. Total ΔG and TΔS show that the most conformational stabilization and order at the base C22.ΔG and TΔS of η show destabilization and disorder at loop1 for holo form with respect to apo **(Fig S10 (c) and (d))**. Holo form with respect to apo form contributes maximum destabilization and disorder at base A21. ΔG and TΔS of η and θ contributes significant stabilization and order at U23. Conformational free energy and entropy for sugar pucker angles corresponding to loop1 of v0, v1, v2, v3, and v4 **(Fig S10 (e) and (f))** do not contribute significantly to the bases. The total changes in conformational thermodynamics due to sugar-puckers show that the most conformational stabilization and order at the base C22 while maximum destabilization and disorder at A20.

#### Loop2

Sugar-phosphate backbone torsion angles along with sugar-base torsion angle corresponding to loop2 do not contribute significantly to bases **(Fig S11. (a) and (b))**. Total changes in conformational free energy and entropy show destabilization and disorder at base A31. Next, we shed light on pseudo-torsion angle **(Fig S11. (c) and (d))**. Destabilization of loop2 is reflected from ΔG of η and θ for (i) holo form with respect to apo. TΔS of □ and θ show significant destabilization and disorder at base A31 and A32. TΔS of θ exhibit the highest stabilization and order at the base A30 for holo form with respect to apo. We find that for sugar-puckers, base A31 has maximum destabilization and disorder **(Fig S11. (e) and (f))**.

#### Ion recognition site

Now we examine the conformational thermodynamics due to sugar-phosphate, sugar-base, and sugar-pucker torsion angles at all bases corresponding to the ion recognition site of Fluoride riboswitch aptamer. OP1/OP2 of P_6_, P_7_, P_8_, P_41_ and P_42_ of RNA participate in coordination to Mg^2+^ (1, 3, 4). Mg^2+^ (2) is coordinated with OP2 of P_40_ while for Mg^2+^ (5), the coordinating residue is O2’ of P_52_. K^+^ is coordinated with OP1 of P_5_, P_6_, P_7_. The rest of the coordination in all the cases is fulfilled by water molecules. Sugar base torsion angle along with sugar-phosphate torsion angles shows ordering and stabilization at all bases corresponding to the ion recognition site **(Fig S12 (a) and (b))**. The total changes in conformational free energy and entropy indicate the maximum value at A6 and U41. We now discuss the changes in conformational thermodynamics in terms of pseudo torsion angles **(Fig S12 (c) and (d))**. We observe that ΔG and TΔS of η and □ at A6, 7U, and A40 corresponding to holo form with respect to apo form show significant stabilization and order while the highest stabilization ΔG and ordering (TΔS) □ at U41 and for η it is G42. Next, we shed light on the conformational thermodynamics for sugar pucker angles **(Fig S12 (e) and (f))**. v0, v1 and v4 exhibit maximum contribution to stabilize the A40 observed from ΔG value. ΔG of v2 and v3 contribute the most at U41. TΔS of v0, v1, v2, v3 and v4 have maximum ordering at U41. Computation of the total changes in conformational free energy and conformational entropy over all sugar-puckers show that G8 has the maximum destabilization and disorder while most stabilization and order are observed at A6 and U41.

**Table 1** shows significant stabilization in pseudoknot; stem1, and ion recognition site of the holo form with respect to the apo form. The intra-base pair is the primary factor to destabilize the holo system compared to the apo form while the inter-base pair, torsion, pseudo-torsion, and sugar angle degrees of freedom are the main factors to stabilize the holo form over the apo form in the pseudoknot region. We find that most of the changes in conformational free energy and entropy are insignificant compared to the thermal energy (≈2.5 kJ/mol) at room temperature for the intra-base pair of pseudoknot. The intra-base pair is also the main factor to destabilize stem2 whereas pseudo-torsion angle and inter-base pairs are the key factors to stabilize the stem1 of the holo system. We observe that loop1 and stem2 get significantly destabilized. We find that most of the changes for loop2 in conformational free energy and entropy (contribution from torsion angle, pseudo-torsion angle sugar angle) are insignificant compared to the thermal energy (≈2.5 kJ/mol) at room temperature.

**Table 1(a).** The changes in conformational thermodynamics of the Pseudoknot region of the holo system in comparison to the apo system (kJ/mol).

| Pseudoknot | Inter Base-pair | Intra Base-pair | Torsion-angle | Pseudotorsion-angle | Sugar-angle | Total |
| --- | --- | --- | --- | --- | --- | --- |
| $\Delta G$(KJ/mol ) | -3.39 | 2.13 | -9.81 | -6.01 | -7.56 | -24.64 |
| $T\Delta S$ (KJ/mol ) | -9.05 | 0.45 | -12.29 | -6.59 | -7.77 | -35.25 |

**Table 1(b).** The changes in conformational thermodynamics of the Stem1 region of the holo system in comparison to the apo system (kJ/mol).

| Stem1 | Inter Base-pair | Intra Base-pair | Torsion-angle | Pseudotorsion-angle | Sugar-angle | Total |
| --- | --- | --- | --- | --- | --- | --- |
| $\Delta G$(KJ/mol ) | -7.57 | -1.53 | -0.75 | -8.68 | -4.26 | -22.79 |
| $T\Delta S$ (KJ/mol ) | -9.98 | -5.73 | -0.44 | -10.45 | -1.14 | -27.74 |

**Table 1(c).** The changes in conformational thermodynamics of the Stem2 region of the holo system in comparison to the apo system (kJ/mol).

| Stem2 | InterBase-pair | IntraBase-pair | Torsion-angle | Pseudotorsion-angle | Sugar-angle | Total |
| --- | --- | --- | --- | --- | --- | --- |
| $\Delta G$(KJ/mol ) | 0.92 | 5.46 | 1.80 | -0.14 | 2.73 | 10.77 |
| $T\Delta S$ (KJ/mol ) | 4.04 | 7.57 | 1.12 | -0.29 | -0.39 | 12.05 |

**Table 1(d).** The changes in conformational thermodynamics of the Loop1 region of the holo system in comparison to the apo system (kJ/mol).

| Loop1 | Torsion-angle | Pseudotorsion-angle | Sugar-angle | Total |
| --- | --- | --- | --- | --- |
| $\Delta G$(KJ/mol ) | 14.13 | 4.05 | 3.14 | 21.32 |
| $T\Delta S$ (KJ/mol ) | 23.74 | 5.51 | 2.47 | 31.72 |

**Table 1(e).**
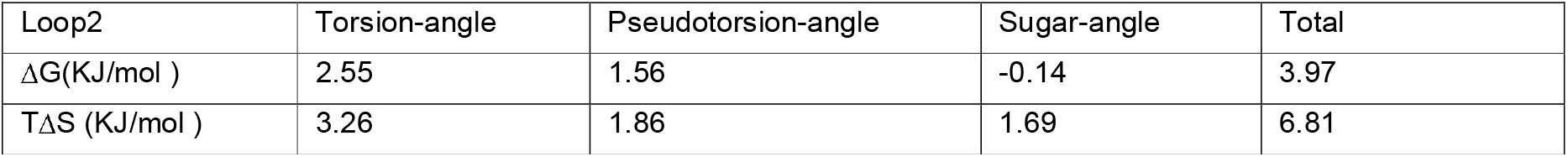
The changes in conformational thermodynamics of the Loop2 region of the holo system in comparison to the apo system (kJ/mol).

**Table 1(f).** The changes in conformational thermodynamics of the Ion recognition site region of the holo system in comparison to the apo system (kJ/mol).

| Ion recognition site | Torsion-angle | Pseudotorsion-angle | Sugar-angle | Total |
| --- | --- | --- | --- | --- |
| $\Delta G$(KJ/mol ) | -23.01 | -20.85 | -15.77 | -59.63 |
| $T\Delta S$ (KJ/mol ) | -30.09 | -23.39 | -17.67 | -71.15 |

We show the stable and flexible regions of the holo system with respect to the apo form in a color-coded cartoon presentation in **Fig S6**.

## 4. Discussion

The health system suffers from a serious crisis in the current century to combat against life-threatening bacterial infections due to the severe emergence of bacterial resistance to antibiotics. Consequently, a new generation of antibiotics with novel mechanisms of action against resistant bacteria is increasingly demanded **[26]**. An artificial cell-based sensor has been developed recently where the encapsulation of Fluoride riboswitch occurs within the lipid vesicles **[27]**. The antibacterial activity of peptides Gramicidin D and Magainin 2 is potentiated in the presence of F^-^ **[28]**. It might be effective to use a combination therapy that consists of proven antibiotic and toxic anion like F^-^ which has antibacterial function.

We discuss the binding mode and interaction pattern of two known antibiotics, Gramicidin D and Magainin 2 peptides towards Fluoride Riboswitch. We further analyze to decipher whether the residues of holo Fluoride riboswitch having instability in conformational free energy and disordering can dock Gramicidin D and Magainin 2. The docked structures are shown in **Fig 4 (a)** for Gramicidin D and **Fig 4 (b)** for Magainin 2.

**Fig. 4:**
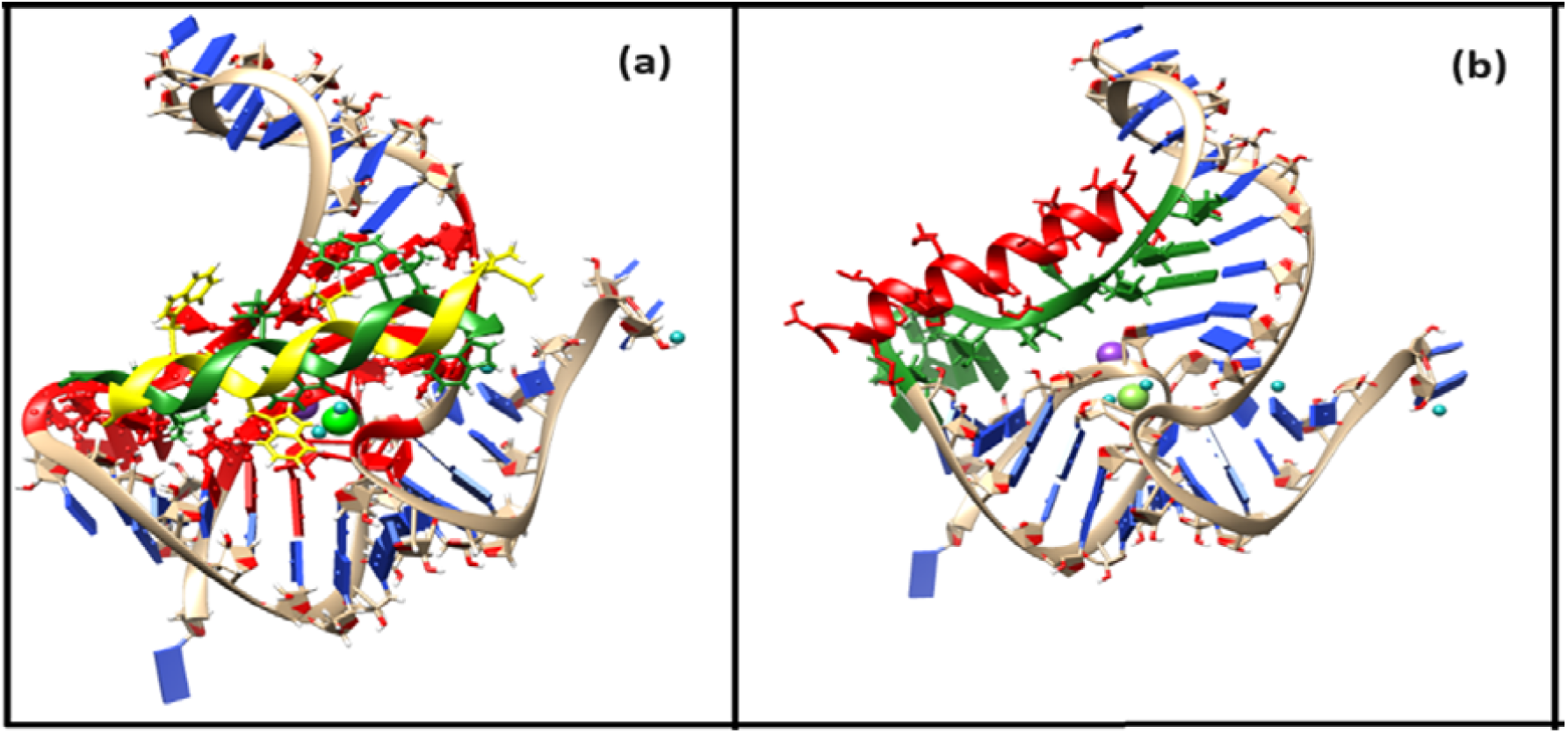
Docked complex of (a) double-stranded (each helix in green and yellow) GramidinD and Fluoride riboswitch (holo) (b) Magainin 2 (red) and Fluoride riboswitch (holo). Binding sites in the receptor Fluoride riboswitch are depicted in (a) red, (b) green

### Gramicidin D binding

Gramicidin D is a linear pentadeca-peptide antibiotic with alternating L and D amino acids. Residues G5, G15, C16, C17, U38 of Pseudoknot, G24, C25, C26, C27, C37 of Stem2, A 21, 22C, 23U of Loop1, 5G, 6A, 7U, 41U, 42G of Ion Recognition Site of holo Fluoride riboswitch act as receptor interface for peptide antibiotic Gramicidin D which is confirmed from conformational thermodynamics data and HDOCK result. Later we perform a biased docking analysis with HADDOCK considering disordered residues A21, C22, U23, G24, C25, C26, C27 as the active residues of Fluoride riboswitch. Residues FVA1, DVA6, Val7, Trp11, DLE12, Trp13 of chain A and Ala3, DLE4, DVA8, Trp9, DLE10, Trp13, DLE14, Trp15 of chain B of Gramicidin D which are found to lie in the vicinity (^L^4.5Å) act as ligand interface for receptor Fluoride riboswitch. Interfacial residues from Gramicidin D show significant hydrophobic interaction.

### Magainin2 binding

The Hebrew word “magain” means “shield”. Conformational thermodynamics data and HDOCK result ensure that destabilized and disordered residues C18, A19, A20, A21, C22, U23 of loop1, 24G, 25C, 26C, 27C, 28U, A33, G34, G35, G36, C37 of stem2 of holo Fluoride riboswitch act as receptor interface. A biased docking analysis with HADDOCK has been performed with Fluoride riboswitch and Magainin 2 considering these destabilized and disordered residues of loop1 and stem2 as the active residues of Fluoride riboswitch. Residues Lys4, Phe5, Ser8, Lys11, Phe12, Lys14, Ala15, Phe16, Gly18, Glu19, Ile20, Met21, Asn22, Ser23 of Magainin 2 which are found to lie in the vicinity (<SUP>L</SUP>4.5Å) act as ligand interface for receptor Fluoride riboswitch. Interfacial residues Ala, Gly, Ile, Met, and Phe from Magainin 2 show significant hydrophobic interaction whereas Lys, and Ser undergo dominant electrostatic interaction with Fluoride riboswitch.

Docking score from HDOCK **(Table S5 (a)) as well as** HADDOCK **(Table S5 (b))** demonstrate that *the Docked complex of Gramidin D* has stronger binding affinity for *Fluoride riboswitch* than that of Magainin 2. HADDOCK result illustrates Van der Waals energy as well as electrostatic energy favour the peptide-RNA complex formation while desolvation energy has less impact on it. Fig. **S13** presents the zoomed view of the interface of the energy-minimized docked complexes. We further dock the peptide drugs Gramicidin D, and Magainin 2, to the average simulated structure of apo Fluoride riboswitch. We show the energy values upon minimization (**Fig. S14 (a), and (b)**) for the docked complexes. The minimum energy for apo Fluoride riboswitch-Gramicidin D complex and apo Fluoride riboswitch-Magainin 2 complex are considerably higher than holo Fluoride riboswitch-Gramicidin D complex and holo Fluoride riboswitch-Magainin 2 complex. Thus we find that peptide drug binding to holo Fluoride riboswitch is more favourable compared to apo Fluoride riboswitch.

## Conclusion

In this study, we investigated the structural, and thermodynamic basis of the interaction of the putative cognate ligand binding with Fluoride riboswitch at the molecular level. Conformational stability and order of the holo form of Fluoride riboswitch aptamer with respect to the apo form is evaluated region wise elaborately by conformational thermodynamic analysis using the histograms of the microscopic conformational variables. The following conclusions emerged from the above study.

Holo Fluoride riboswitch gets energetically and entropically stable at the Pseudoknot, Stem1, and Ion recognition sites. However, Loop1, Loop2, Stem2, and most of the unpaired bases show significant disorder and destabilization. Pseudoknot-like reversed Watson-Crick A6•U38 base pairing and reversed Hoogsteen A40•U48 base pairing play a critical role in stabilizing Fluoride riboswitch, one of the higher-order RNA architectures. Thermodynamically destabilized and disordered residues from Loop1 and Stem2 serve as putative binding sites for non-cognate ligands. In summary, these findings provide new insight into RNA-ligand interaction. Exploring the possible role of nucleotides in the ligand binding site of Fluoride riboswitch aptamer may enable the design of new ligands and aptamers with the development of nucleic acid-based therapeutics.

## Author Contributions

Soumi Das conceptualized, curated, analyzed, and interpreted the data and wrote the manuscript.

## Acknowledgments

Soumi Das is thankful to the Technical Research Centre, S. N. Bose National Centre for Basic Sciences, Kolkata for the computational facilities, and the Department of Science and Technology (DST) for funding.

