## Supplementary Information for "In-silico studies on thermodynamics of ligand binding to Fluoride riboswitch aptamer"

**Author: Soumi Das^1*^**

**^1^Department of Physics of Complex Systems, S. N. Bose National Centre for Basic Sciences, Block-JD, Sector-III, Salt Lake, Kolkata 700106, India.**

**Correspondence: Soumi Das**

**List of all the microscopic conformational variables of RNA used in our study are described:**

i. inter-bp step parameters: tilt (τ), roll (ρ), twist (ω), shift (D_x_), slide (D_y_), and rise (D_z_).
ii. intra-bp step parameters: buckle (κ), open (σ), propeller (π), stagger (S_x_), shear (S_y_), stretch (S_z_).
iii. sugar-phosphate backbone torsion angles: α, β, γ, δ, ε, ζ and sugar-base backbone torsion angle χ.
iv. pseudo torsion angle (ƞ and θ).
v. sugar pucker: (ν_0_, ν_1_, ν_2_, ν_3_, ν_4_)

**The Nomenclature and Description of the RNA structural parameters:**

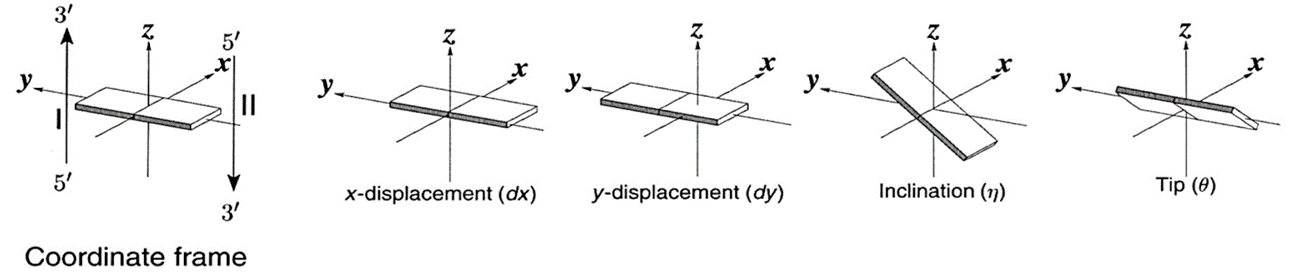

**RNA inter-base pair parameters [1]**

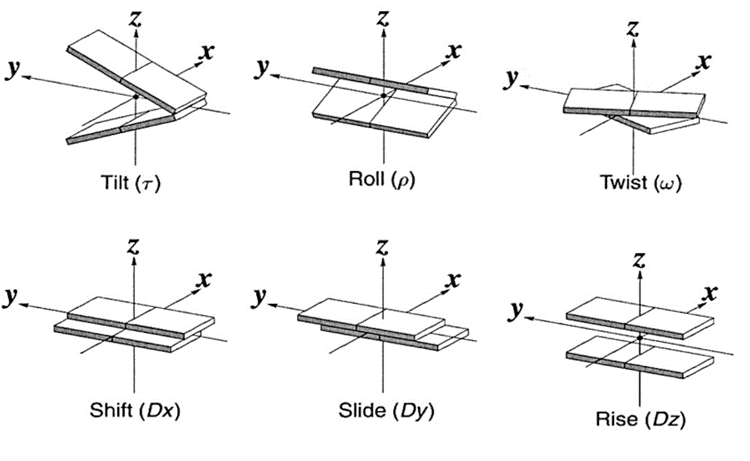

**RNA intra-base pair parameters [2]**

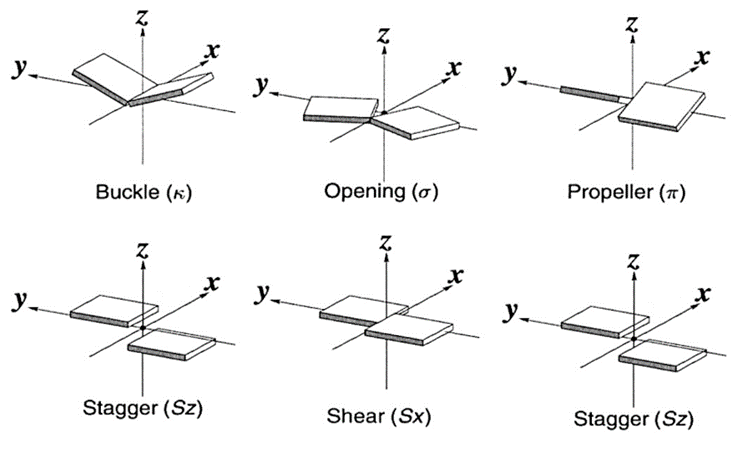

**RNA Sugar-phosphate torsion angle and Sugar-pucker (endocyclic torsion angle) [3]**

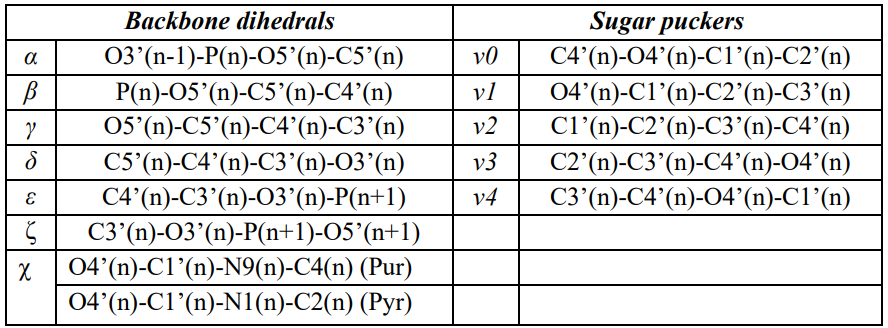

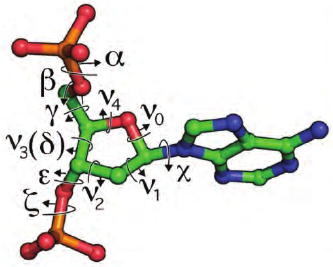

**RNA Pseudotorsion angle [4]**

A virtual bond between P and C4’ is introduced for defining pseudo-torsion angles η (C4'_N-1_-P_N_-C4'_N_-P_N+1_) and θ (P_N_-C4'_N_-P_N+1_-C4'_N+1_) in order to describe reduced dimensionality of RNA backbone configuration **[4]**.

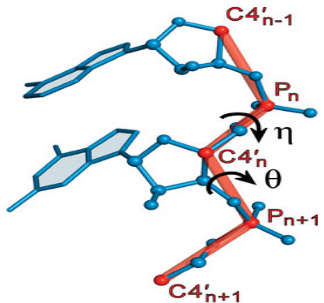

**RNA Sugar-base torsion angle [3]**

Dihedral χ is defined around the glycosidic bonds C1’–N9 for purines and C1’–N1 for pyrimidines [3]. This dihedral represents the rotation of the nucleobase with respect to the sugar ring. Anti (χ ~ 180˚ ± 90˚) and syn (χ ~ 0˚ ± 90˚) conformations are defined based on the torsion angle of the glycosidic bond. The sequence of atoms chosen to define anti/syn conformation is: O4’-C1’-N9-C4 for purines and O4’-C1’-N1-C2 for pyrimidines.

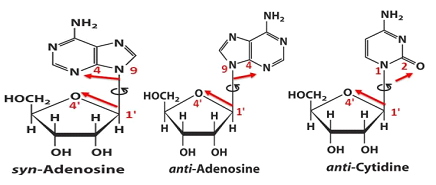

**RNA** **Pseudo-rotation phase angle (P) [3]**

Sugar geometry is often represented by pseudo-rotation phase angle P and amplitude of puckering ν_max_, where P and ν_max_ are given by the following expression:

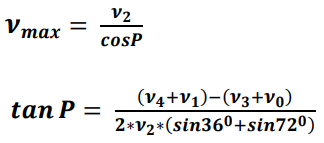

All the endocyclic torsion angles are related to P and ν_max_ by the following equation:

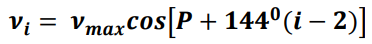

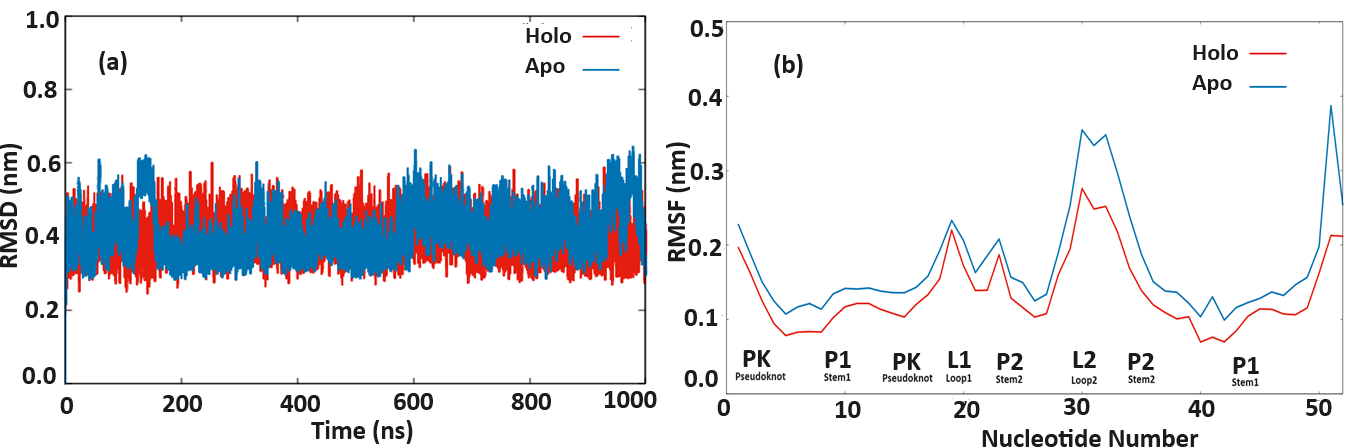

**Fig. S1 (a): Root mean square deviation (RMSD) plots of backbone atoms of Holo (red) and Apo (blue) Fluoride riboswitch aptamer as a function of simulation time. (b): RMSF around average nucleotides position of 1 µs trajectories of Holo (red) and Apo (blue) riboswitches.**

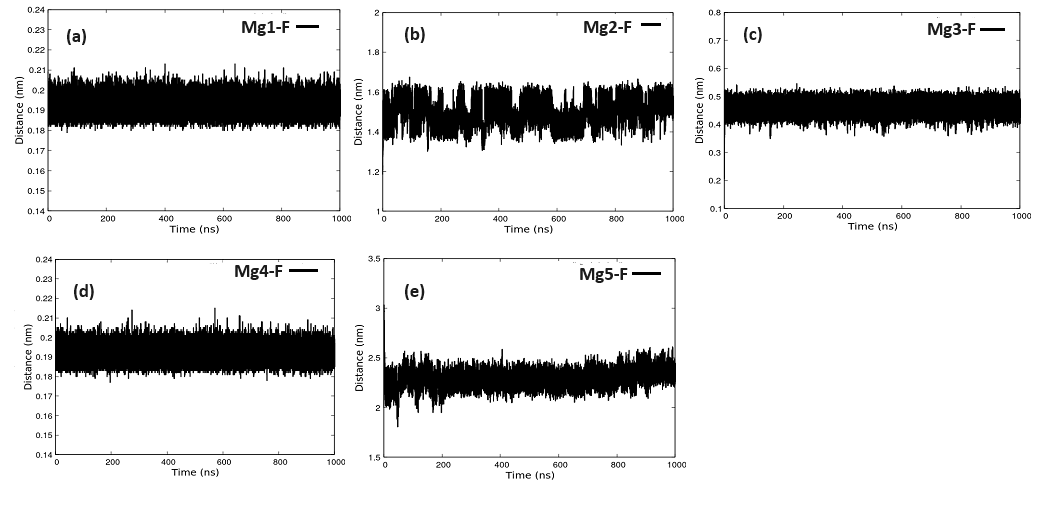

**Fig. S2 (a) distance between Mg1 and F, (b) distance between Mg2 and F, (c) distance between Mg3 and F, (d) distance between Mg4 and F, and (f) distance between Mg5 and F**

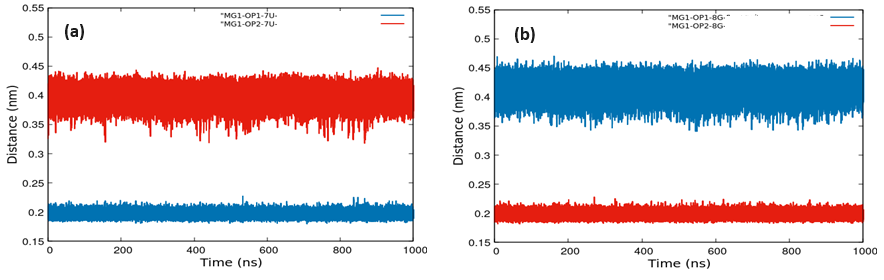

**Fig. S3 (a) distance between Mg1 and OP1/OP2-7U, and (b) distance between Mg1 and OP1/OP2-8G**

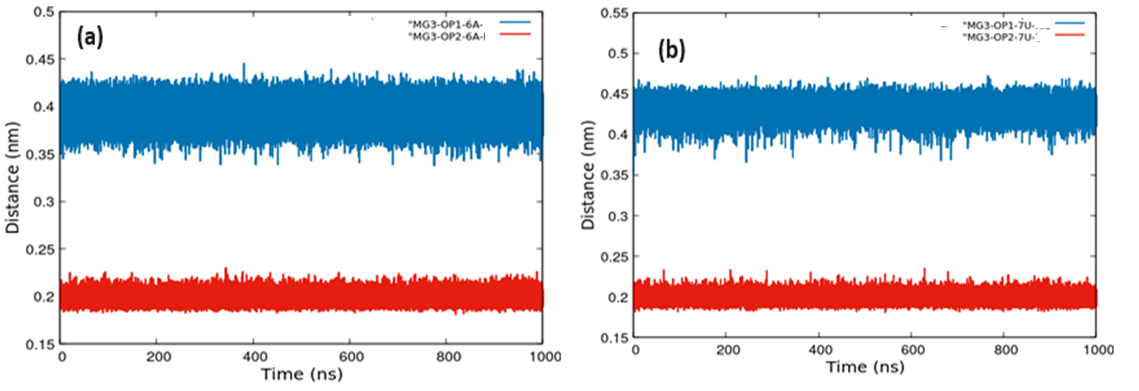

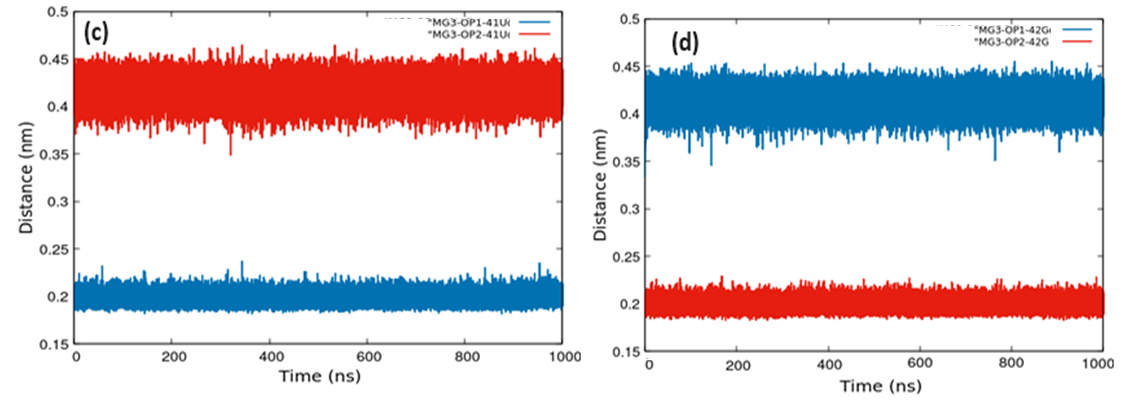

**Fig. S4 (a) distance between Mg3 and OP1/OP2-6A, (b) distance between Mg3 and OP1/OP2-7U,**

**(c) distance between Mg3 and OP1/OP2-41U, and (d) distance between Mg3 and OP1/OP2-42G**

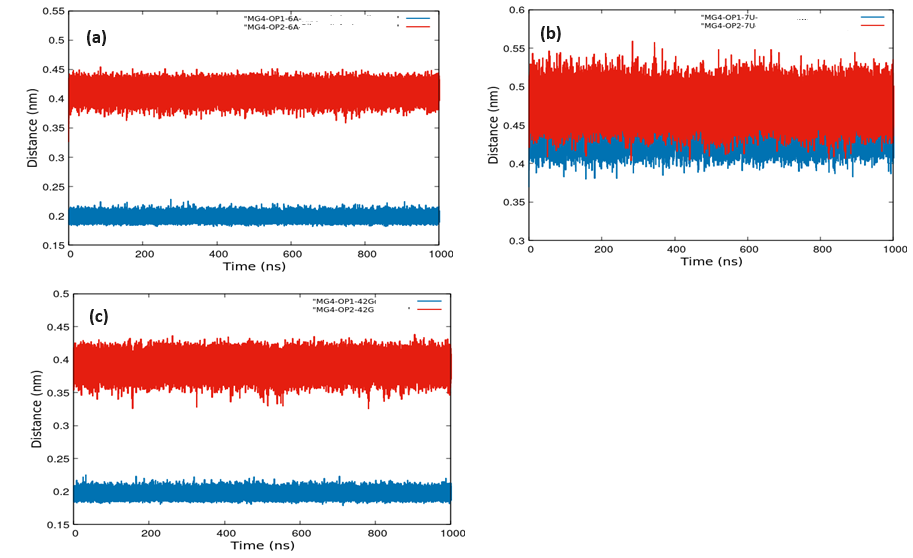

**Fig. S5 (a) distance between Mg4 and OP1/OP2-6A, (b) distance between Mg4 and OP1/OP2-7U, and (c) distance between Mg4 and OP1/OP2-42G**

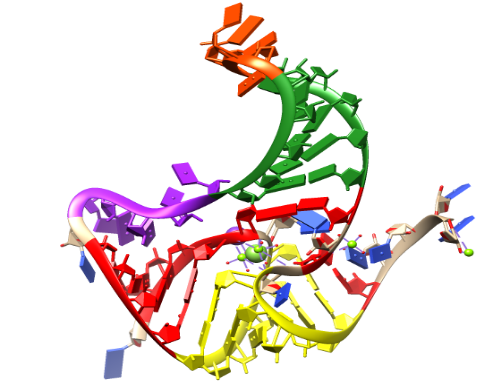

**Fig S6: Colour coding of residue wise conformational thermodynamics holo system of fluoride riboswitch with respect to the apo form:**

**red: Pseudoknot	(highly stable and ordered),**

**yellow: Stem1	(highly stable and ordered),**

**green: Stem2	(highly unstable and disordered),**

**purple: Loop1	(highly unstable and disordered),**

**orange: Loop2	(highly unstable and disordered)**

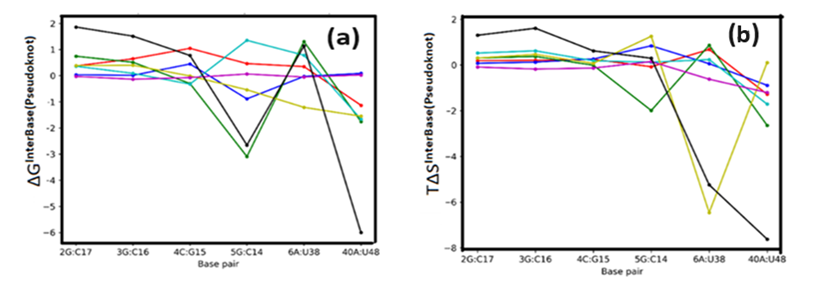

**Fig S7: ΔG (a) and TΔS (b): Pseudoknot due to inter-bp step parameters ("Tilt" in red, "Roll" in blue, "Twist" in green, "Shift" in cyan, "Slide" in yellow, "Rise" in magenta, "Total" in black)**

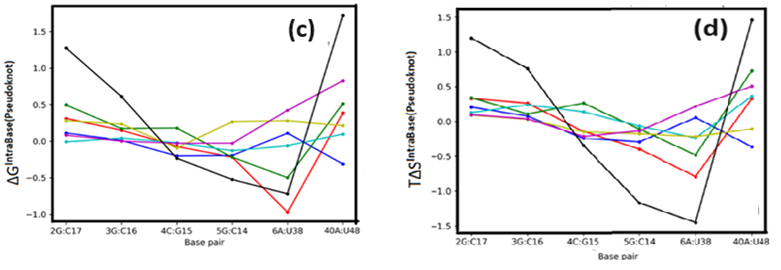

**Fig S7: ΔG (c) and TΔS (d): Pseudoknot due to intra-bp step parameters ("Buckle" in red, "Open" in blue, "Propeller" in green, "Shear" in cyan, "Stagger" in yellow, "Stretch" in magenta, "Total" in black)**

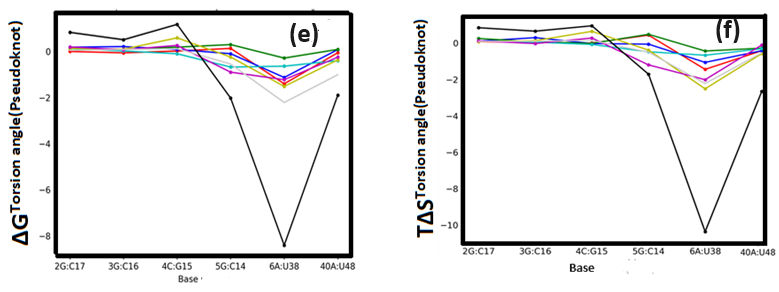

**Fig S7: ΔG (e) and TΔS (f): Pseudoknot due to Torsion Angle ("alpha" in red, "beta" in blue, "gamma" in green, "delta" in cyan, "eps" in yellow, "zeta" in magenta, "chi" in light gray, "Total" in black)**

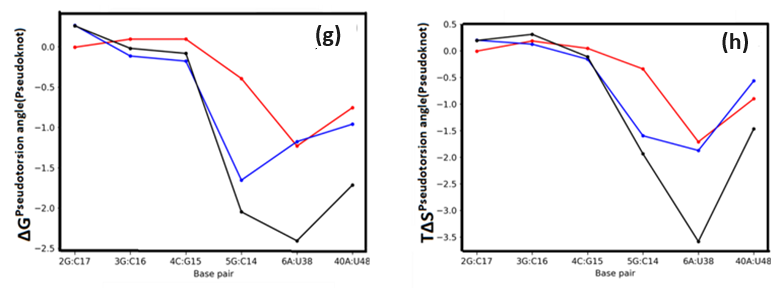

**Fig S7: ΔG (g) and TΔS (h): Pseudoknot due to Pseudotorsion Angle ("Eta" in red, "Theta" in blue, "Total" in black)**

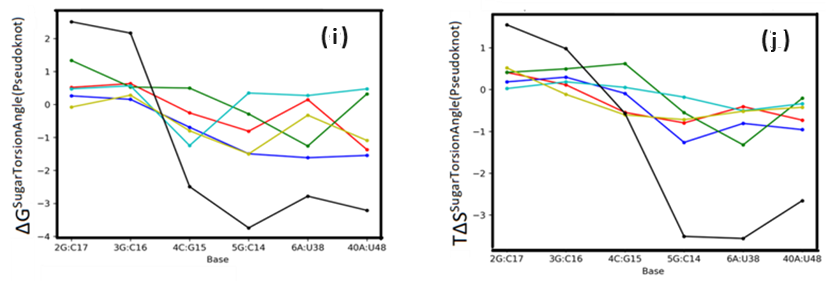

**Fig S7: ΔG (i) and TΔS (j): Pseudoknot due to Sugar Torsion Angle (ν0 in red, ν1 in blue, ν2 in green, ν3 in cyan, ν4 in yellow, "Total" in black)**

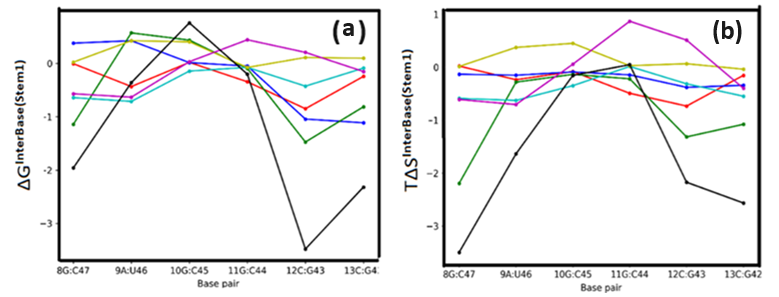

**Fig S8: ΔG (a) and TΔS (b): Stem1 due to inter-bp step parameters ("Tilt" in red, "Roll" in blue, "Twist" in green, "Shift" in cyan, "Slide" in yellow, "Rise" in magenta, "Total" in black)**

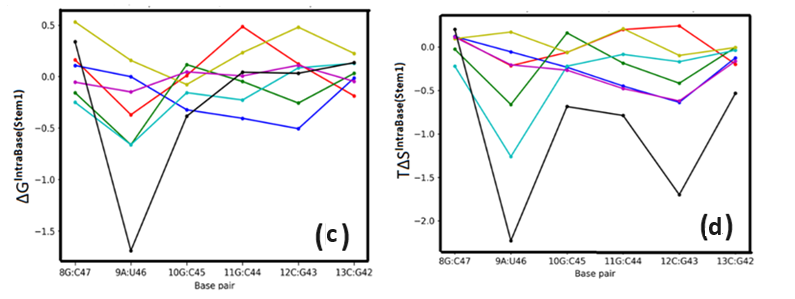

**Fig S8: ΔG (c) and TΔS (d): Stem1 due to intra-bp step parameters ("Buckle" in red, "Open" in blue, "Propeller" in green, "Shear" in cyan, "Stagger" in yellow, "Stretch" in magenta, "Total" in black)**

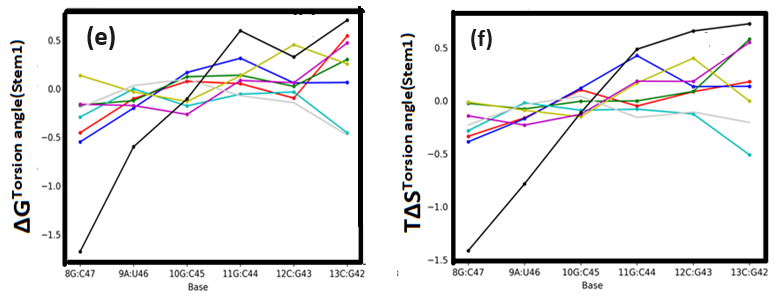

**Fig S8: ΔG (e) and TΔS (f): Stem1 due to Torsion Angle ("alpha" in red,"beta" in blue, "gamma" in green, "delta" in cyan, "eps" in yellow, "zeta" in magenta, "chi" in light gray, "Total" in black)**

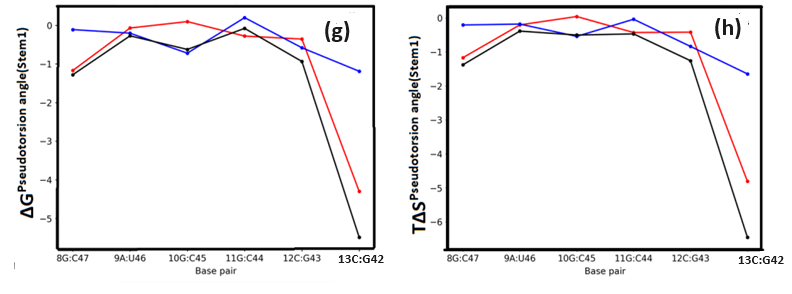

**Fig S8: ΔG (g) and TΔS (h): Stem1 due to due to Pseudotorsion Angle ("Eta" in red, "Theta" in blue, "Total" in black)**

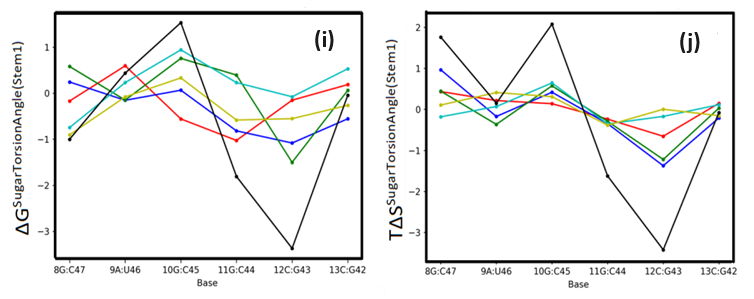

**Fig S8: ΔG (i) and TΔS (j): Stem1 due to Sugar Torsion Angle (ν0 in red, ν1 in blue, ν2 in green, ν3 in cyan, ν4 in yellow, "Total" in black)**

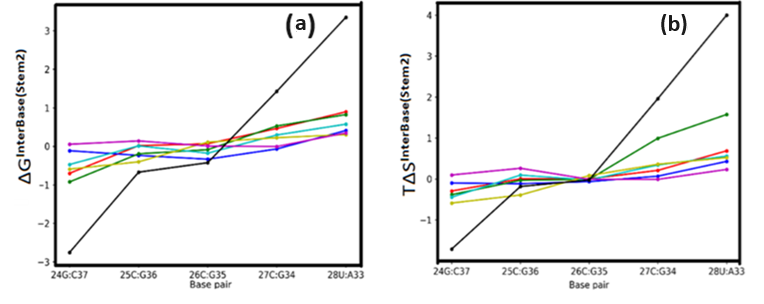

**Fig S9: ΔG (a) and TΔS (b): Stem2 due to inter-bp step parameters ("Tilt" in red, "Roll" in blue, "Twist" in green, "Shift" in cyan, "Slide" in yellow, "Rise" in magenta, "Total" in black)**

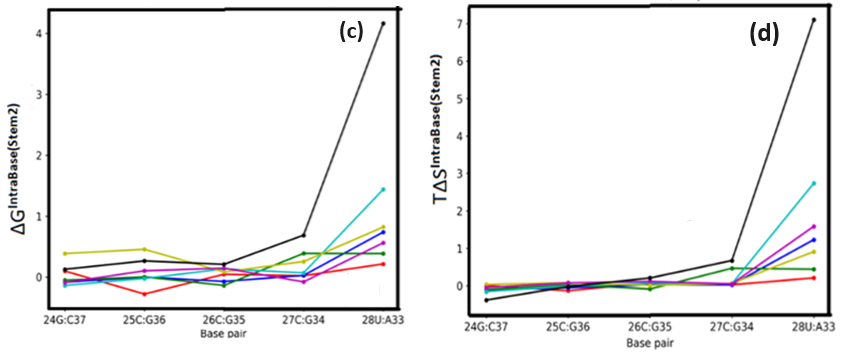

**Fig S9: ΔG (c) and TΔS (d): Stem2 due to intra-bp step parameters ("Buckle" in red, "Open" in blue, "Propeller" in green, "Shear" in cyan, "Stagger" in yellow, "Stretch" in magenta, "Total" in black)**

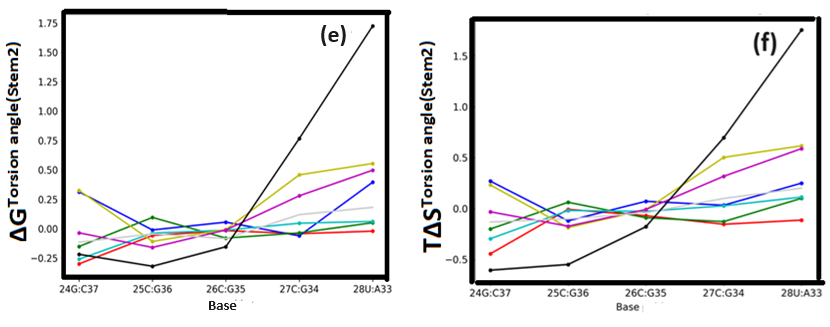

**Fig S9: ΔG (e) and TΔS (f): Stem2 due to Torsion Angle ("alpha" in red, "beta" in blue, "gamma" in green, "delta" in cyan, "eps" in yellow, "zeta" in magenta, "chi" in light grey, "Total" in black)**

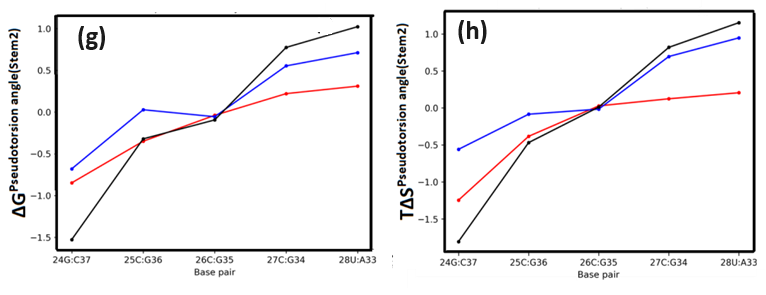

**Fig S9: ΔG (g) and TΔS (h): Stem2 due to due to Pseudotorsion Angle ("Eta" in red, "Theta" in blue, "Total" in black)**

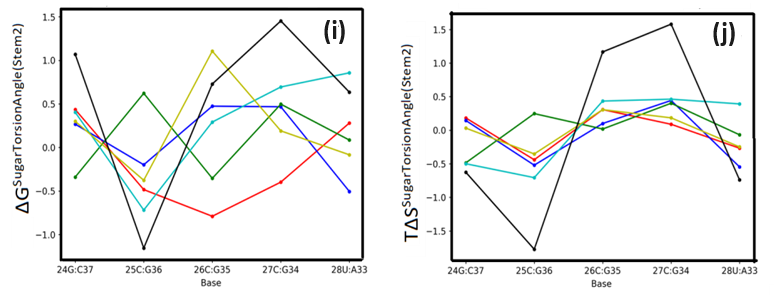

**Fig S9: ΔG (i) and TΔS (j): Stem2 due to Sugar Torsion Angle (ν0 in red, ν1 in blue, ν2 in green, ν3 in cyan, ν4 in yellow, "Total" in black)**

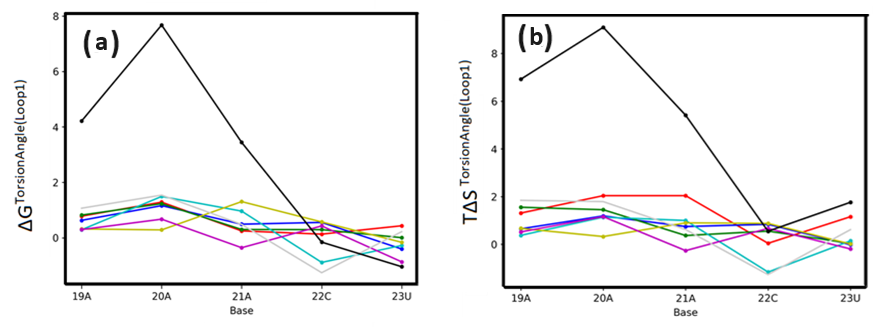

**Fig S10: ΔG (a) and TΔS (b): Loop1 due to Torsion Angle ("alpha" in red, "beta" in blue, "gamma" in green, "delta" in cyan, "eps" in yellow, "zeta" in magenta, "chi" in light grey, "Total" in black)**

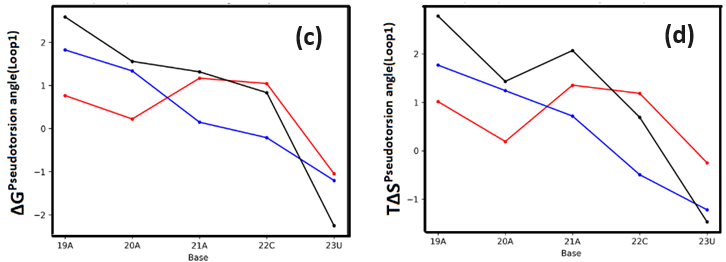

**Fig S10: ΔG (c) and TΔS (d): Loop1 due to Pseudotorsion Angle ("Eta" in red, "Theta" in blue, "Total" in black)**

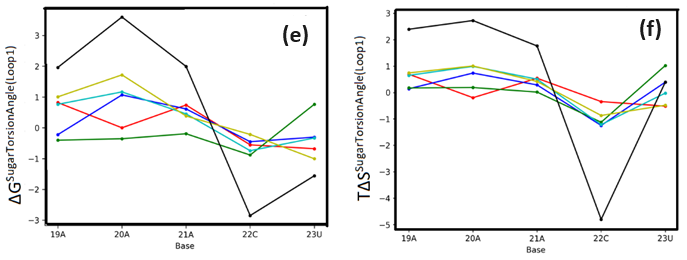

**Fig S10: ΔG (e) and TΔS (f): Loop1 due to Sugar Torsion Angle (ν0 in red, ν1 in blue, ν2 in green, ν3 in cyan, ν4 in yellow, "Total" in black)**

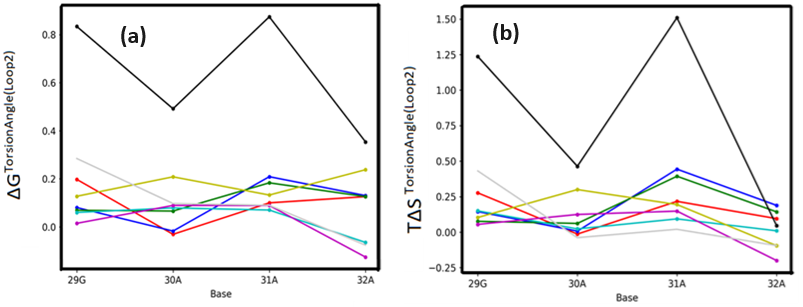

**Fig S11: ΔG (a) and TΔS (b): Loop2 due to Torsion Angle ("alpha" in red, "beta" in blue, "gamma" in green, "delta" in cyan, "eps" in yellow, "zeta" in magenta, "chi" in light grey, "Total" in black)**

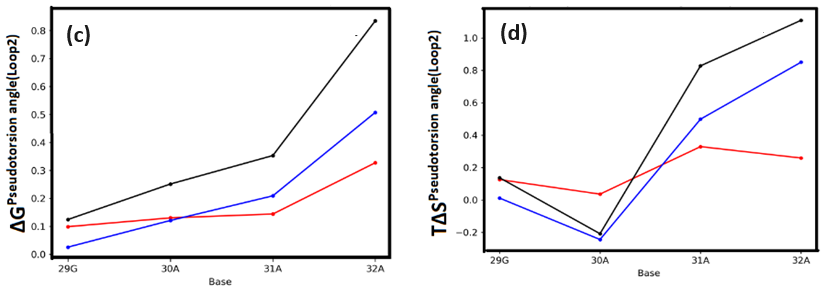

**Fig S11: ΔG (c) and TΔS (d): Loop2 due to Pseudotorsion Angle ("Eta" in red, "Theta" in blue, "Total" in black)**

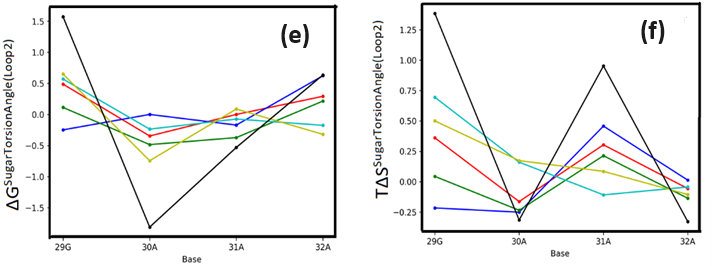

**Fig S11: ΔG (e) and TΔS (f): Loop2 due to Sugar Torsion Angle (ν0 in red, ν1 in blue, ν2 in green, ν3 in cyan, ν4 in yellow, "Total" in black)**

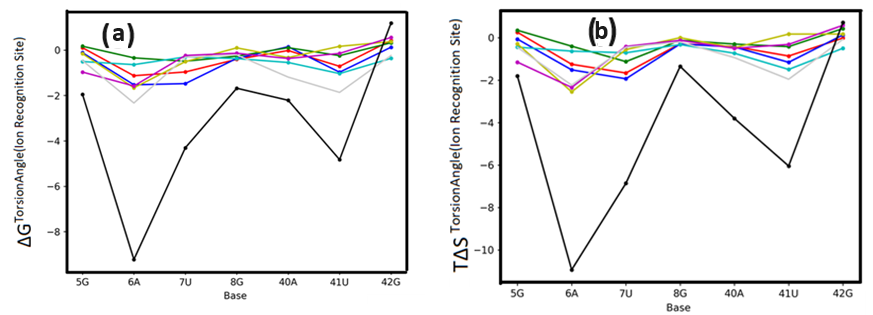

**Fig S12: ΔG (a) and TΔS (b): Ion Recognition Site due to Torsion Angle ("alpha" in red, "beta" in blue, "gamma" in green, "delta" in cyan, "eps" in yellow, "zeta" in magenta, "chi" in light grey, "Total" in black)**

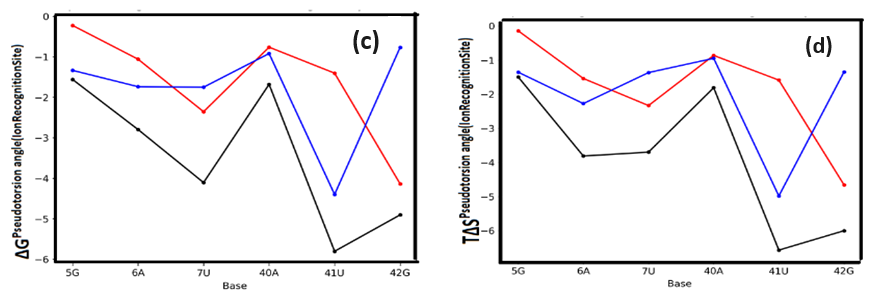

**Fig S12: ΔG (c) and TΔS (d): Ion Recognition Site due to Pseudotorsion Angle ("Eta" in red, "Theta" in blue, "Total" in black)**

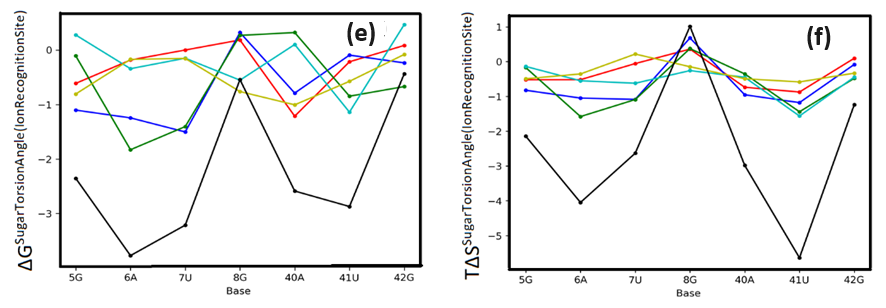

**Fig S12: ΔG (e) and TΔS (f): Ion recognition Site due to Sugar Torsion Angle (ν0 in red, ν1 in blue, ν2 in green, ν3 in cyan, ν4 in yellow, "Total" in black)**

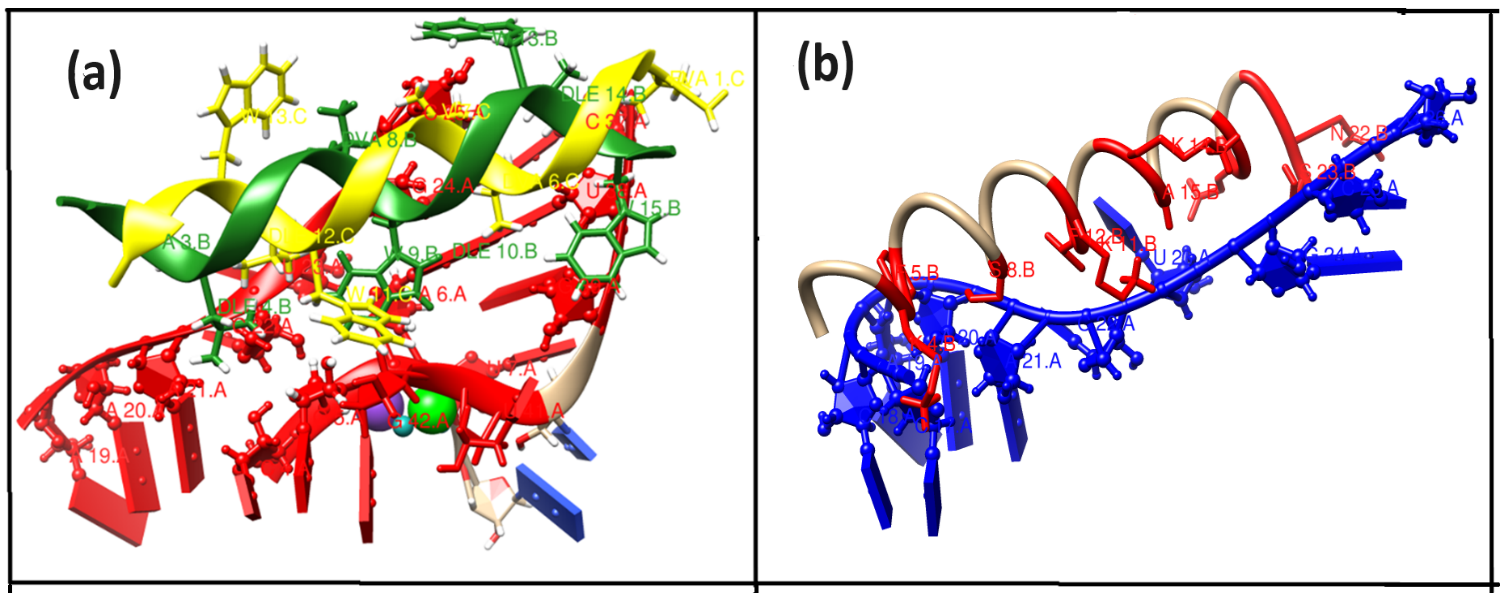

***Fig. S13: Zoomed view of the interface of the energy-minimized docked complex (a) GramidinD-Fluoride riboswitch (holo) (b) Magainin 2 and Fluoride riboswitch (holo).***

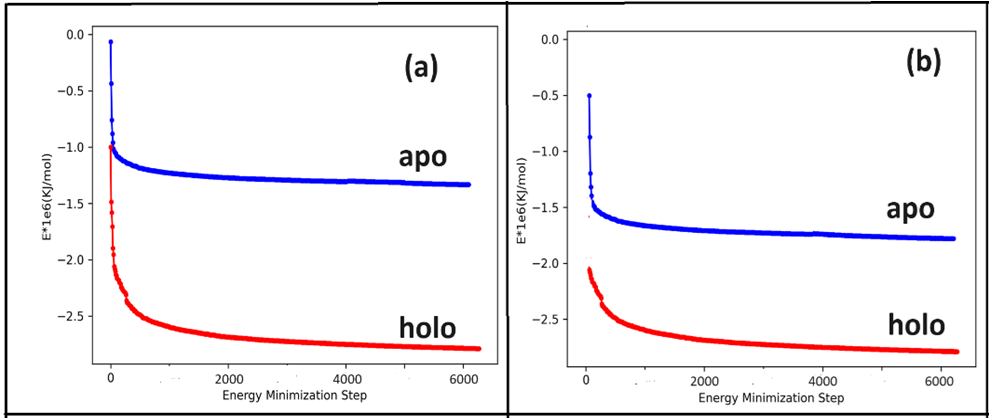

***Fig. S14 The energy-minimization plots corresponding to (a) Gramicidin D (b) Magainin 2 complexes with apo (blue) and holo (red) Fluoride riboswitch***

**Table S1 (a): RMSD (Mean and SD) of region wise riboswitch in nm**

| Region | System | RMSD (mean and SD) in nm |
| --- | --- | --- |
| Pseudoknot | Holo | 0.21 (0.04) |
|  | Apo | 0.19 (0.04) |
| Stem1 | Holo | 0.09 (0.01) |
|  | Apo | 0.13 (0.02) |
| Stem2 | Holo | 0.11 (0.02) |
|  | Apo | 0.11 (0.02) |
| Loop1 | Holo | 0.10 (0.02) |
|  | Apo | 0.13 (0.02) |
| Loop2 | Holo | 0.08 (0.02) |
|  | Apo | 0.07 (0.01) |
| Ligand Binding Site | Holo | 0.12 (0.04) |
|  | Apo | 0.34 (0.03) |

**Table S1 (b): % of Hydrogen bond occupancy within base pairs constituting different region of the aptamer domain.**

|  |  |  | Occupancy (%) | |
| --- | --- | --- | --- | --- |
| Region | Base Pairing | donor-acceptor | Holo | Apo |
| Pseudoknot | 2 G:C 17 (W:W C) | 2G (N2-H21) - 17 C (O2 ) | 99.9 | 29.9 |
|  |  | 2G (N1-H1) - 17 C (N3 ) | 99.9 | 15.9 |
|  | 3 G:C 16 (W:W C) | 3G (N1-H1 ) - 16 C (N3 ) | 99.8 | 27.3 |
|  |  | 3G (N2-H21) - 16 C (O2 ) | 99.9 | 12.9 |
|  | 4 C:G 15 (W:W C) | 4C (N4-H41) - 15 G (O6 ) | 94.0 | 21.3 |
|  |  | 15G (N2-H21) - 4 C (O2 ) | 99.6 | 19.6 |
|  |  | 15G (N1-H1) - 4 C (N3 ) | 99.8 | 29.8 |
|  | 5 G:C 14 (W:W C) | 5G (N1-H1 ) - 14 C (N3 ) | 99.9 | 10.9 |
|  |  | 5G (N2-H21) - 14 C (O2 ) | 99.5 | 26.7 |
|  | 6 A:U 38 (W:W T) | 6A (N6-H61) - 38 U (O2 ) | 98.1 | 25.1 |
|  |  | 38U (N3-H3) - 6 A (N1 ) | 99.1 | 21.9 |
|  | 40 A:U 48  (H:W T) | 40A (N6-H61) - 48 U (O2 ) | 99.6 | 23.5 |
|  |  | 48U (N3-H3) - 40 A (N7 ) | 98.9 | 22.3 |
| Stem 1 | 8 G:C 47 (W:W C) | 8G (N1-H1) - 47 C (N3 ) | 99.9 | 24.9 |
|  |  | 8G (N2-H21) - 47 C (O2 ) | 99.8 | 22.7 |
|  | 9 A:U 46 (W:W C) | 46U (N3-H3) - 9 A (N1 ) | 98.9 | 23.1 |
|  |  | 9A (N6-H61) - 46 U (O4 ) | 87.8 | 26.7 |
|  | 10 G:C 45 (W:W C) | 10G (N1-H1) - 45 C (N3 ) | 99.9 | 23.9 |
|  |  | 10G (N2-H21) - 45 C (O2 ) | 99.6 | 23.5 |
|  | 11 G:C 44 (W:W C) | 11G (N1-H1) - 44 C (N3 ) | 99.9 | 21.8 |
|  |  | 11G (N2-H21) - 44 C (O2 ) | 99.7 | 22.7 |
|  | 12 C:G 43 (W:W C) | 12C (N4-H41) - 43 G (O6 ) | 96.6 | 24.1 |
|  |  | 43G (N2-H21) - 12 C (O2 ) | 99.8 | 26.7 |
|  |  | 43G (N1-H1 ) - 12 C (N3 ) | 99.8 | 28.2 |
|  | 13 C:G 42 (W:W C) | 42G (N2-H21) - 13 C (O2 ) | 99.9 | 27.8 |
|  |  | 42G (N1-H1) - 13 C (N3 ) | 99.9 | 24.9 |
| Stem 2 | 24 G:C 37 (W:W C) | 24G (N1-H1) - 37 C (N3 ) | **69.9** | **19.8** |
|  |  | 24G (N2-H21) - 37 C (O2 ) | **69.8** | **19.9** |
|  |  | 37C (N4-H41) - 24 G (O6 ) | **78.8** | **28.4** |
|  | 25 C:G 36 (W:W C) | 25C (N4-H41) - 36 G (O6 ) | **78.1** | **28.2** |
|  |  | 36G (N2-H21) - 25 C (O2 ) | **59.8** | **19.9** |
|  |  | 36G (N1-H1 ) - 25 C (N3 ) | **49.8** | **21.8** |
|  | 26 C:G 35 (W:W C) | 26C (N4-H41) - 35 G (O6 ) | **78.3** | **28.3** |
|  |  | 35G (N2-H21) - 26 C (O2 ) | **79.8** | **23.9** |
|  |  | 35G (N1-H1 ) - 26 C (N3 ) | **79.7** | **29.8** |
|  | 27 C:G 34 (W:W C) | 27C (N4-H41) - 34 G (O6 ) | **77.8** | **27.8** |
|  |  | 34G (N2-H21) - 27 C (O2 ) | **79.7** | **29.8** |
|  |  | 34G (N1-H1 ) - 27 C (N3 ) | **79.8** | **29.8** |
|  | 28 U:A 33 (W:W C) | 28U (N3-H3 ) - 33 A (N1 ) | **77.9** | **28.8** |
|  |  | 33A (N6-H61) - 28 U (O4 ) | **77.7** | **28.5** |

**Table S1(c): Bond distance (nm) between the ions (ligand) and the ligand binding nucleotides in the holoriboswitch**

| Bond distance | MD simulation (nm) | X-ray (nm) |
| --- | --- | --- |
| **Mg1-F** | 0.19(0.003) | 0.19 |
| **Mg2-F** | 1.50(0.06) | 1.17 |
| **Mg3-F** | 0.46(0.02) | 0.18 |
| **Mg4-F** | 0.19(0.003) | 0.19 |
| **Mg5-F** | 2.30(0.08) | 2.55 |
| Mg1-OP1-7U | 0.19(0.004) | 0.21 |
| Mg1-OP2-7U | 0.39(0.01) | 0.34 |
| Mg1-OP1-8G | 0.42(0.01) | 0.42 |
| Mg1-OP2-8G | 0.19(0.004) | 0.19 |
| Mg3-OP1-6A | 0.39(0.01) | 0.36 |
| Mg3-OP2-6A | 0.19(0.005) | 0.19 |
| Mg3-OP1-7U | 0.43(0.009) | 0.36 |
| Mg3-OP2-7U | 0.19(0.005) | 0.21 |
| Mg3-OP1-41U | 0.19(0.005) | 0.19 |
| Mg3-OP2-41U | 0.41(0.01) | 0.43 |
| Mg3-OP1-42G | 0.41(0.01) | 0.32 |
| Mg3-OP2-42G | 0.19(0.005) | 0.20 |
| Mg4-OP1-6A | 0.19(0.004) | 0.20 |
| Mg4-OP2-6A | 0.41(0.01) | 0.33 |
| Mg4-OP1-7U | 0.44(0.01) | 0.35 |
| Mg4-OP2-7U | 0.48(0.02) | 0.42 |
| Mg4-OP1-42G | 0.20(0.004) | 0.20 |
| Mg4-OP2-42G | 0.39(0.01) | 0.37 |
| K-OP1-5G | **2.28(1.17)** | **0.25** |
| K-OP2-5G | **2.28(1.19)** | **0.39** |
| K-OP1-6A | **2.27(1.18)** | **0.28** |
| K-OP2-6A | **2.29(1.16)** | **0.43** |
| K-OP1-7U | **2.29(1.18)** | **0.23** |
| K-OP2-7U | **2.31(1.17)** | **0.44** |

**Table S2 (a): Mean and SD of Inter Base pair parameter (holo)**

| **Base pairing** | **Tilt** | **Roll** | **Twist** | **Shift** | **Slide** | **Rise** |
| --- | --- | --- | --- | --- | --- | --- |
| **Pseudoknot** |  |  |  |  |  |  |
| **2 G:C 17 (W:W C)** | **0.83(2.59)** | **6.77(4.57)** | **28.94(2.75)** | **0.06(0.35)** | **-2.12(0.31)** | **3.23(0.22)** |
| **3 G:C 16 (W:W C)** | **0.33(2.45)** | **2.99(3.95)** | **31.71(2.82)** | **-0.24(0.37)** | **-1.87(0.41)** | **3.23(0.21)** |
| **4 C:G 15 (W:W C)** | **-0.16(2.40)** | **9.30(6.05)** | **31.24(2.65)** | **-0.27(0.43)** | **-1.31(0.36)** | **3.40(0.22)** |
| **5 G:C 14 (W:W C)** | **3.98(7.38)** | **11.01(6.49)** | **19.58(4.33)** | **-1.21(0.43)** | **0.36(0.42)** | **-1.16(3.56)** |
| **6 A:U 38 (W:W T)** | **-6.02(7.30)** | **-6.68(8.5)** | **-22.01(9.21)** | **-3.48(0.67)** | **-3.57(0.75)** | **6.49(0.32)** |
| **40 A:U 48 (H:W T)** | **-1.74(5.11)** | **8.83(7.40)** | **36.37(8.49)** | **-1.80(0.80)** | **-3.18(0.50)** | **3.41(0.3)** |
| **Stem1** |  |  |  |  |  |  |
| **8 G:C 47 (W:W C)** | **-1.47(2.63)** | **10.73(5.16)** | **30.37(3.04)** | **-0.38(0.42)** | **-1.92(0.34)** | **3.28(0.24)** |
| **9 A:U 46 (W:W C)** | **-1.71(2.75)** | **12.56(5.01)** | **28.93(3.22)** | **-0.61(0.47)** | **-1.55(0.38)** | **3.22(0.23)** |
| **10 G:C 45 (W:W C)** | **0.32(2.43)** | **14.37(4.89)** | **29.71(2.74)** | **0.10(0.39)** | **-1.57(0.31)** | **3.24(0.21)** |
| **11 G:C 44 (W:W C)** | **2.47(2.33)** | **6.31(4.14)** | **31.92(2.77)** | **0.35(0.35)** | **-1.39(0.39)** | **3.27(0.22)** |
| **12 C:G 43 (W:W C)** | **3.18(2.51)** | **5.36(4.18)** | **34.50(2.60)** | **0.74(0.32)** | **-1.48(0.34)** | **3.53(0.19)** |
| **13 C:G 42 (W:W C)** | **-0.42(2.25)** | **3.17(4.46)** | **27.30(2.60)** | **-0.43(0.32)** | **-1.80(0.30)** | **3.40(0.22)** |
| **Stem2** |  |  |  |  |  |  |
| **24 G:C 37 (W:W C)** | **0.023(2.59)** | **2.44(4.14)** | **32.06(3.54)** | **-0.13(0.37)** | **-2.13(0.42)** | **3.23(0.23)** |
| **25 C:G 36 (W:W C)** | **0.589(2.53)** | **7.16(4.85)** | **29.41(2.96)** | **-0.04(0.39)** | **-2.02(0.33)** | **3.31(0.23)** |
| **26 C:G 35 (W:W C)** | **0.18(2.53)** | **7.86(5.12)** | **29.67(3.13)** | **-0.05(0.41)** | **-1.94(0.34)** | **3.61(0.27)** |
| **27 C:G 34 (W:W C)** | **-0.90(2.93)** | **7.62(5.95)** | **25.30(6.42)** | **-0.15(0.53)** | **-1.61(0.41)** | **3.06(0.33)** |
| **28 U:A 33 (W:W C)** | **11.98(7.8)** | **17.01(9.27)** | **44.80(6.35)** | **1.08(0.66)** | **-0.83(0.49)** | **1.93(1.88)** |
| **Loop1** |  |  |  |  |  |  |
| **18 C:** | **6.12(30.78)** | **-23.56(57.64)** | **-27.19(18.27)** | **-8.07(2.24)** | **-2.36(1.85)** | **0.88(3.27)** |
| **19 A:** | **10.88(110.25)** | **12.90(51.92)** | **-2.63(17.37)** | **1.95(2.50)** | **3.18(1.17)** | **4.73(0.59)** |
| **20 A:Cs:s C** | **2.41(6.60)** | **19.84(6.72)** | **3.72(5.69)** | **-0.64(0.44)** | **0.88(0.53)** | **5.42(0.90)** |
| **21 A:G 3 (s:s T)** | **-38.59(12.71)** | **47.07(19.39)** | **50.82(14.54)** | **3.04(1.07)** | **3.84(0.82)** | **2.69(1.61)** |
| **22 C:** | **-12.69(41.60)** | **-7.49(60.3)** | **15.03(23.18)** | **0.03(2.08)** | **-0.35(1.3)** | **1.55(3.95)** |
| **23 U:** | **26.25(77.71)** | **-32.04(64.76)** | **-28.05(30.92)** | **-10.47(3.59)** | **8.91(6.8)** | **3.28(0.22)** |
| **Loop2** |  |  |  |  |  |  |
| **29 G:** | **81.49(65.9)** | **-52.86(50.61)** | **21.62(21.79)** | **-4.92(1.06)** | **6.49(1.12)** | **3.21(0.32)** |
| **30 A:** | **9.82(6.72)** | **-3.0(11.5)** | **37.85(15.01)** | **1.01(1.29)** | **-1.47(0.75)** | **3.30(0.28)** |
| **31 A:** | **5.90(6.81)** | **-2.63(8.85)** | **33.51(10.33)** | **0.06(1.06)** | **-1.59(0.64)** | **2.11(0.75)** |
| **32 A:** | **16.2 (9.11)** | **10.17(13.89)** | **76.41(9.14)** | **8.09(0.88)** | **-1.24(0.69)** | **3.61(0.27)** |
| **Ion recognition-site** |  |  |  |  |  |  |
| **5 G:C 14 (W:W C)** | **7.77(49.85)** | **-124.38(42.73)** | **-43.39(25.49)** | **-12.03(4.34)** | **15.51(7.16)** | **6.85(4.95)** |
| **6 A:U 38 (W:W T)** | **-6.02(7.30)** | **-6.68(8.5)** | **-22.01(9.21)** | **-3.48(0.67)** | **-3.57(0.75)** | **6.49(0.32)** |
| **7 U:** | **-76.78(7.94)** | **4.5(8.4)** | **43.16(8.96)** | **-1.07(0.73)** | **-2.98(0.65)** | **9.60(0.50)** |
| **8 G:C 47 (W:W C)** | **-1.47(2.63)** | **10.73(5.16)** | **30.37(3.04)** | **-0.38(0.42)** | **-1.92(0.34)** | **3.28(0.24)** |
| **40 A:U 48 (H:W T)** | **-1.74(5.11)** | **8.83(7.40)** | **36.37(8.49)** | **-1.80(0.80)** | **-3.18(0.50)** | **3.41(0.3)** |
| **41 U:Cs:h T** | **10.00(4.05)** | **-6.74(5.74)** | **-6.71(4.23)** | **-3.65(0.52)** | **-3.5(0.43)** | **3.27(0.22)** |
| **42 G:C 13 (W:W C)** | **-3.18(2.51)** | **5.36(4.18)** | **34.50(2.60)** | **-0.74(0.32)** | **-1.48(0.34)** | **3.24(0.21)** |

**Table S2 (b): Mean and SD of Intra Base pair parameter (holo)**

| **Base pairing** | **Buckle** | **Open** | **Propeller** | **Stagger** | **Shear** | **Stretch** |
| --- | --- | --- | --- | --- | --- | --- |
| **Pseudoknot** |  |  |  |  |  |  |
| **2 G:C 17 (W:W C)** | **-3.66(8.02)** | **0.35(3.28)** | **-5.9(7.4)** | **-0.15(0.29)** | **-0.15(0.26)** | **2.91(0.08)** |
| **3 G:C 16 (W:W C)** | **-8.07(8.65)** | **0.20(3.29)** | **-11.30(6.35)** | **-0.07(0.31)** | **-0.002(0.32)** | **2.91(0.09)** |
| **4 C:G 15 (W:W C)** | **-6.19(7.87)** | **-0.63(3.36)** | **-12.23(6.93)** | **0.01(0.3)** | **-0.02(0.31)** | **2.91(0.08)** |
| **5 G:C 14 (W:W C)** | **-2.89(7.36)** | **-2.23(3.07)** | **-1.70(6.39)** | **0.04(0.29)** | **-0.04(0.29)** | **2.93(0.08)** |
| **6 A:U 38 (W:W T)** | **-2.95(8.47)** | **-1.06(5.79)** | **-6.90(7.85)** | **-0.20(0.37)** | **-0.17(0.27)** | **2.93(0.13)** |
| **40 A:U 48 (H:W T)** | **-5.05(9.1)** | **4.87(5.32)** | **-12.20(7.37)** | **-0.08(0.39)** | **-0.07(0.28)** | **2.91(0.12)** |
| **Stem1** |  |  |  |  |  |  |
| **8 G:C 47 (W:W C)** | **-3.36(8.48)** | **-0.78(3.30)** | **-9.23(6.89)** | **0.01(0.3)** | **0.04(0.28)** | **2.93(0.09)** |
| **9 A:U 46 (W:W C)** | **-5.09(8.69)** | **4.56(6.10)** | **-14.36(7.77)** | **-0.16(0.41)** | **0.04(0.27)** | **2.89(0.11)** |
| **10 G:C 45 (W:W C)** | **-5.02(8.25)** | **-0.99(3.35)** | **-11.08(7.42)** | **-0.19(0.31)** | **-0.006(0.30)** | **2.90(0.08)** |
| **11 G:C 44 (W:W C)** | **-7.27(9.25)** | **-0.09(3.29)** | **-12.21(6.50)** | **-0.18(0.31)** | **-0.02(0.29)** | **2.90(0.08)** |
| **12 C:G 43 (W:W C)** | **-3.24(9.38)** | **-0.10(3.48)** | **-13.14(6.19)** | **-0.13(0.31)** | **0.08(0.30)** | **2.90(0.08)** |
| **13 C:G 42 (W:W C)** | **-4.59(7.30)** | **-1.15(3.22)** | **-9.68(6.43)** | **-0.004(0.3)** | **0.14(0.30)** | **2.91(0.08)** |
| **Stem2** |  |  |  |  |  |  |
| **24 G:C 37 (W:W C)** | **2.48(8.68)** | **-1.20(3.17)** | **-4.36(6.78)** | **-0.01(0.31)** | **-0.11(0.29)** | **2.92(0.08)** |
| **25 C:G 36 (W:W C)** | **3.37(8.18)** | **0.32(3.45)** | **-8.49(6.60)** | **-0.14(0.31)** | **0.05(0.30)** | **2.91(0.09)** |
| **26 C:G 35 (W:W C)** | **0.96(8.09)** | **0.005(3.54)** | **-7.89(6.58)** | **-0.17(0.29)** | **0.17(0.31)** | **2.91(0.09)** |
| **27 C:G 34 (W:W C)** | **0.55(8.15)** | **-0.53(3.44)** | **-4.63(7.91)** | **-0.15(0.31)** | **0.10(0.30)** | **2.92(0.09)** |
| **28 U:A 33 (W:W C)** | **9.16(9.78)** | **-1.17(7.40)** | **3.57(9.83)** | **0.03(0.50)** | **1.05(2.24)** | **2.87(0.29)** |
| **Loop1** |  |  |  |  |  |  |
| **18 C:** | **0.0** | **0.0** | **0.0** | **0.0** | **0.0** | **0.0** |
| **19 A:** | **0.0** | **0.0** | **0.0** | **0.0** | **0.0** | **0.0** |
| **20 A:Cs:s C** |  |  |  |  |  |  |
| **21 A:G 3 (s:s T)** | **-30.41(12.2)** | **-57.78(6.3)** | **-24.1(7.5)** | **1.16(0.5)** | **-2.48(0.3)** | **2.46(0.3)** |
| **22 C:** | **0.0** | **0.0** | **0.0** | **0.0** | **0.0** | **0.0** |
| **23 U:** | **0.0** | **0.0** | **0.0** | **0.0** | **0.0** | **0.0** |
| **Loop2** |  |  |  |  |  |  |
| **29 G:** | **0.0** | **0.0** | **0.0** | **0.0** | **0.0** | **0.0** |
| **30 A:** | **0.0** | **0.0** | **0.0** | **0.0** | **0.0** | **0.0** |
| **31 A:** | **0.0** | **0.0** | **0.0** | **0.0** | **0.0** | **0.0** |
| **32 A:** | **0.0** | **0.0** | **0.0** | **0.0** | **0.0** | **0.0** |
| **Ion recognition-site** |  |  |  |  |  |  |
| **5 G:C 14 (W:W C)** | **-2.89(7.36)** | **-2.23(3.07)** | **-1.70(6.39)** | **0.04(0.29)** | **-0.04(0.29)** | **2.93(0.08)** |
| **6 A:U 38 (W:W T)** | **-2.95(8.47)** | **-1.06(5.79)** | **-6.90(7.85)** | **-0.20(0.37)** | **-0.17(0.27)** | **2.93(0.13)** |
| **7 U:** | **0.0** | **0.0** | **0.0** | **0.0** | **0.0** | **0.0** |
| **8 G:C 47 (W:W C)** | **-3.36(8.48)** | **-0.78(3.30)** | **-9.23(6.89)** | **0.01(0.3)** | **0.04(0.28)** | **2.93(0.09)** |
| **40 A:U 48 (H:W T)** | **-5.05(9.1)** | **4.87(5.32)** | **-12.20(7.37)** | **-0.08(0.39)** | **-0.07(0.28)** | **2.91(0.12)** |
| **41 U:Cs:h T** | **22.32(12.96)** | **39.26(9.13)** | **-5.57(12.10)** | **-0.01(0.86)** | **2.25(0.59)** | **3.98(0.57)** |
| **42 G:C 13 (W:W C)** | **4.59(7.30)** | **-1.15(3.22)** | **-9.68(6.43)** | **-0.004(0.30)** | **-0.15(0.30)** | **2.91(0.09)** |

**Table S2(c): Mean and SD of The torsion and pseudo torsion angles for holo**

| **Base** | **CHI** | **PHASE** | **ETA** | **THETA** |
| --- | --- | --- | --- | --- |
| **Pseudoknot** |  |  |  |  |
| **2G** | **-156.49** \| **(57.32)** | **10.75** \| **(10.01)** | **158.56** \| **(46.33)** | **-148.83** \| **(13.64)** |
| **3G** | **-145.97** \| **(85.41)** | **8.37** \| **(9.91)** | **120.08** \| **(123.24)** | **-146.96** \| **(19.55)** |
| **4C** | **-160.67** \| **(26.69)** | **12.20** \| **(9.43)** | **139.43** \| **(98.78)** | **-151.07** \| **(12.92)** |
| **5G** | **-160.42** \| **(29.23)** | **6.47** \| **(9.48)** | **141.06** \| **(97.09)** | **76.92** \| **(5.02)** |
| **6A** | **-13.98** \| **(173.38)** | **9.31** \| **(11.15)** | **-154.31** \| **(3.84)** | **145.18** \| **(95.92)** |
| **14C** | **-158.04** \| **(16.87)** | **17.60** \| **(10.81)** | **152.69** \| **(72.13)** | **-152.1** \| **(15.64)** |
| **15G** | **-158.63** \| **(31.59)** | **8.82** \| **(10.28)** | **160.18** \| **(37.20)** | **-142.27** \| **(7.86)** |
| **16C** | **-159.64** \| **(23.22)** | **13.63** \| **(10.29)** | **153.68** \| **(67.66)** | **-146.83** \| **(9.51)** |
| **17C** | **-159.91** \| **(23.96)** | **16.09** \| **(10.09)** | **146.85** \| **(84.20)** | **-155.04** \| **(14.24)** |
| **38U** | **-158.41** \| **(15.56)** | **17.05** \| **(10.43)** | **159.40** \| **(42.67)** | **-137.06** \| **(10.68)** |
| **40A** | **-121.53** \| **(9.27)** | **121.27** \| **(14.49)** | **-56.36** \| **(7.83)** | **-112.96** \| **(6.57)** |
| **48U** | **-158.03** \| **(24.67)** | **19.50** \| **(27.50)** | **161.29** \| **(30.56)** | **-146.60** \| **(44.74)** |
| **Stem1** |  |  |  |  |
| **8G** | **-128.77** \| **(111.53)** | **21.04** \| **(12.52)** | **123.23** \| **(118.65)** | **-149.69** \| **(19.00)** |
| **9A** | **-157.55** \| **(18.51)** | **18.16** \| **(13.00)** | **153.35** \| **(62.43)** | **-152.35** \| **(26.51)** |
| **10G** | **-157.96** \| **(22.63)** | **14.30** \| **(10.92)** | **159.58** \| **(28.37)** | **-148.53** \| **(10.64)** |
| **11G** | **-159.27** \| **(32.80)** | **13.28** \| **(11.03)** | **153.93** \| **(66.31)** | **-144.54** \| **(8.58)** |
| **12C** | **-159.74** \| **(28.30)** | **13.49** \| **(10.59)** | **156.85** \| **(54.78)** | **-143.11** \| **(11.87)** |
| **13C** | **-160.34** \| **(20.01)** | **16.14** \| **(11.41)** | **160.06** \| **(47.51)** | **-152.11** \| **(14.40)** |
| **42G** | **35.88** \| **(170.42)** | **-26.57** \| **(12.07)** | **3.68** \| **(5.04)** | **-113.99** \| **(4.72)** |
| **43G** | **-158.2** \| **(23.26)** | **14.06** \| **(15.41)** | **156.80** \| **(8.95)** | **-134.80** \| **(7.04)** |
| **44C** | **-153.68** \| **(9.32)** | **24.48** \| **(13.08)** | **155.36** \| **(7.29)** | **-149.97** \| **(9.74)** |
| **45C** | **-155.45** \| **(9.13)** | **21.37** \| **(10.68)** | **159.39** \| **(29.09)** | **-144.06** \| **(9.32)** |
| **46U** | **-155.77** \| **(11.66)** | **21.93** \| **(11.27)** | **160.29** \| **(27.61)** | **-151.06** \| **(12.69)** |
| **47C** | **-158.71** \| **(21.78)** | **21.79** \| **(11.05)** | **153.29** \| **(68.45)** | **-148.05** \| **(15.34)** |
| **Stem2** |  |  |  |  |
| **24G** | **-28.19** \| **(171.21)** | **-9.77** \| **(14.24)** | **-122.24** \| **(113.44)** | **-124.44** \| **(12.61)** |
| **25C** | **-159.96** \| **(31.50)** | **14.98** \| **(12.59)** | **152.81** \| **(55.71)** | **-145.57** \| **(12.61)** |
| **26C** | **-160.15** \| **(23.76)** | **18.69** \| **(12.38)** | **153.56** \| **(66.20)** | **-148.57** \| **(16.57)** |
| **27C** | **-157.52** \| **(14.39)** | **22.77** \| **(13.13)** | **147.45** \| **(81.42)** | **-147.04** \| **(21.61)** |
| **28U** | **-149.85** \| **(10.97)** | **22.26** \| **(12.09)** | **158.90** \| **(38.50)** | **-139.93** \| **(28.25)** |
| **33A** | **-157.53** \| **(44.67)** | **8.87** \| **(9.51)** | **157.38** \| **(16.20)** | **-143.71** \| **(8.77)** |
| **34G** | **-157.78** \| **(43.65)** | **11.40** \| **(10.39)** | **149.73** \| **(77.99)** | **-149.26** \| **(9.76)** |
| **35G** | **-157.85** \| **(51.39)** | **11.47** \| **(9.97)** | **151.20** \| **(72.46)** | **-149.42** \| **(9.13)** |
| **36G** | **-148.28** \| **(80.75)** | **9.42** \| **(10.43)** | **150.02** \| **(76.91)** | **-144.89** \| **(9.12)** |
| **37C** | **-160.38** \| **(30.25)** | **14.45** \| **(10.44)** | **90.08** \| **(147.73)** | **-143.82** \| **(9.61)** |
| **Loop1** |  |  |  |  |
| **18C** | **-132.98** \| **(10.97)** | **139.35** \| **(60.91)** | **-153.11** \| **(8.33)** | **109.87** \| **(16.25)** |
| **19A** | **-87.99** \| **(110.45)** | **81.61** \| **(142.84)** | **-138.81** \| **(24.25)** | **122.20** \| **(32.13)** |
| **20A** | **-103.35** \| **(24.18)** | **-13.80** \| **(91.30)** | **119.37** \| **(108.07)** | **-9.98** \| **(157.93)** |
| **21A** | **-96.98** \| **(14.00)** | **-2.48** \| **(26.11)** | **-43.85** \| **(161.59)** | **-98.96** \| **(16.29)** |
| **22C** | **-138.61** \| **(15.07)** | **32.16** \| **(27.44)** | **135.45** \| **(76.94)** | **-141.10** \| **(31.28)** |
| **23U** | **-99.46** \| **(79.72)** | **43.92** \| **(46.14)** | **150.91** \| **(35.00)** | **72.72** \| **(21.06)** |
| **Loop2** |  |  |  |  |
| **29G** | **-146.54** \| **(13.09)** | **18.06** \| **(11.78)** | **156.21** \| **(17.63)** | **-125.65** \| **(15.33)** |
| **30A** | **-114.23** \| **(120.75)** | **0.61** \| **(17.66)** | **35.91** \| **(12.49)** | **-124.53** \| **(11.03)** |
| **31A** | **-151.66** \| **(45.70)** | **9.69** \| **(13.04)** | **127.59** \| **(97.29)** | **-117.48** \| **(93.22)** |
| **32A** | **-154.54** \| **(35.83)** | **9.97** \| **(10.12)** | **155.93** \| **(62.84)** | **-100.90** \| **(9.98)** |
| **Ion recognition-site** |  |  |  |  |
| **5G** | **-160.42** \| **(29.23)** | **6.47** \| **(9.48)** | **141.06** \| **(97.09)** | **76.92** \| **(5.02)** |
| **6A** | **-13.98** \| **(173.38)** | **9.31** \| **(11.15)** | **-154.31** \| **(3.84)** | **145.18** \| **(95.92)** |
| **7U** | **-130.59** \| **(9.38)** | **86.94** \| **(12.75)** | **-127.88** \| **(4.25)** | **-143.08** \| **(7.12)** |
| **8G** | **-128.77** \| **(111.53)** | **21.04** \| **(12.52)** | **123.23** \| **(118.65)** | **-149.69** \| **(19.00)** |
| **40A** | **-121.53** \| **(9.27)** | **121.27** \| **(14.49)** | **-56.36** \| **(7.83)** | **-112.96** \| **(6.57)** |
| **41U** | **-153.24** \| **(51.08)** | **13.77** \| **(16.59)** | **-162.99** \| **(8.61)** | **-90.60** \| **(4.16)** |
| **42G** | **35.88** \| **(170.42)** | **-26.57** \| **(12.07)** | **3.68** \| **(5.04)** | **-113.99** \| **(4.72)** |

**Table S3 (a): Mean and SD of Inter Base parameter (apo)**

| **Base pairing** | **Tilt** | **Roll** | **Twist** | **Shift** | **Slide** | **Rise** |
| --- | --- | --- | --- | --- | --- | --- |
| **Pseudoknot** |  |  |  |  |  |  |
| **2 G:C 17 (W:W C)** | **1.32(2.53)** | **7.37(4.35)** | **28.29(2.51)** | **0.11(0.28)** | **-2.16(0.30)** | **3.20(0.21)** |
| **3 G:C 16 (W:W C)** | **0.92(2.51)** | **3.15(3.85)** | **32.06(2.67)** | **-0.06(0.34)** | **-1.93(0.35)** | **3.19(0.20)** |
| **4 C:G 15 (W:W C)** | **1.33(2.62)** | **8.37(5.51)** | **30.49(2.82)** | **-0.08(0.42)** | **-1.43(0.40)** | **3.52(0.24)** |
| **5 G:C 14 (W:W C)** | **5.23(7.19)** | **11.47(6.42)** | **18.50(4.17)** | **-1.34(0.43)** | **0.42(0.39)** | **3.16(0.29)** |
| **6 A:U 38 (W:W T)** | **-10.97(8.81)** | **11.29(9.66)** | **-24.32(9.27)** | **-4.25(0.97)** | **-1.53(0.87)** | **6.62(0.50)** |
| **40 A:U 48 (H:W T)** | **-56.43(59.3)** | **33.32(14.62)** | **15.52(16.38)** | **5.17(1.20)** | **-0.47(4.10)** | **8.53(2.74)** |
| **Stem1** |  |  |  |  |  |  |
| **8 G:C 47 (W:W C)** | **-1.52(2.78)** | **13.06(5.49)** | **32.81(3.82)** | **-0.30(0.47)** | **-1.93(0.32)** | **3.13(0.24)** |
| **9 A:U 46 (W:W C)** | **-0.92(2.91)** | **13.04(5.14)** | **28.72(3.92)** | **-0.29(0.56)** | **-1.66(0.35)** | **3.23(0.23)** |
| **10 G:C 45 (W:W C)** | **0.97(2.47)** | **15.26(5.01)** | **28.53(2.99)** | **0.36(0.39)** | **-1.77(0.28)** | **3.26(0.22)** |
| **11 G:C 44 (W:W C)** | **0.80(2.34)** | **5.84(4.38)** | **30.50(3.11)** | **0.53(0.36)** | **-1.53(0.43)** | **3.46(0.22)** |
| **12 C:G 43 (W:W C)** | **2.71(2.47)** | **9.74(4.54)** | **28.67(3.27)** | **0.23(0.37)** | **-1.91(0.31)** | **3.23(0.22)** |
| **13 C:G 42 (W:W C)** | **0.27(2.45)** | **7.22(4.59)** | **26.55(3.11)** | **-0.26(0.37)** | **-1.99(0.34)** | **3.36(0.21)** |
| **Stem2** |  |  |  |  |  |  |
| **24 G:C 37 (W:W C)** | **-0.03(2.53)** | **2.26(4.24)** | **31.80(3.51)** | **-0.24(0.36)** | **-1.91(0.50)** | **3.34(0.22)** |
| **25 C:G 36 (W:W C)** | **0.61(2.54)** | **7.88(5.00)** | **29.63(2.94)** | **0.01(0.38)** | **-1.95(0.32)** | **3.25(0.23)** |
| **26 C:G 35 (W:W C)** | **0.26(2.55)** | **8.67(5.08)** | **29.56(3.07)** | **-0.01(0.42)** | **-1.92(0.33)** | **3.29(0.23)** |
| **27 C:G 34 (W:W C)** | **-1.60(2.62)** | **7.26(5.79)** | **28.10(3.24)** | **-0.29(0.45)** | **-1.73(0.36)** | **3.60(0.26)** |
| **28 U:A 33 (W:W C)** | **9.83(5.88)** | **14.27(7.81)** | **42.63(4.60)** | **1.01(0.61)** | **-0.89(0.44)** | **3.13(0.29)** |
| **Loop1** |  |  |  |  |  |  |
| **18 C:** | **13.43(14.46)** | **-50.76(18.93)** | **-34.1(10.13)** | **-9.14(0.81)** | **-2.58(0.82)** | **-0.17(1.49)** |
| **19 A:** | **-50.8 (29.13)** | **-104.58(57.74)** | **26.16(18.73)** | **6.02(1.93)** | **7.37(1.25)** | **1.64(1.29)** |
| **20 A:Cs:s C** | **11.82(3.35)** | **4.36(4.70)** | **68.22(3.31)** | **2.29(0.32)** | **-0.91(0.28)** | **3.42(0.27)** |
| **21 A:G 3 (s:s T)** | **-44.1 (17.07)** | **44.66(21.56)** | **61.98(25.69)** | **3.35(1.88)** | **4.34(1.13 )** | **5.73(1.43)** |
| **22 C:** | **19.63(17.05)** | **0.45(15.87)** | **10.90(20.60)** | **-2.31(2.21)** | **-1.26(1.19)** | **3.17(0.87)** |
| **23 U:** | **76.38(60.08)** | **-56.01(34.49)** | **-33.8(27.09)** | **-8.98(3.93)** | **9.28(5.90)** | **4.77(2.37)** |
| **Loop2** |  |  |  |  |  |  |
| **29 G:** | **81.41(66.57)** | **-53.02(49.77)** | **20.24(20.58)** | **-4.82(1.06)** | **6.43(1.09)** | **2.10(1.83)** |
| **30 A:** | **9.13(7.37)** | **-2.22(12.24)** | **40.30(13.44)** | **1.21(1.18)** | **-1.40(0.86)** | **3.24(0.36)** |
| **31 A:** | **6.96(6.50)** | **-3.32(8.63)** | **31.63(9.21)** | **-0.06(0.90)** | **-1.50(0.69)** | **3.27(0.27)** |
| **32 A:** | **16.74(8.71)** | **11.68(13.86)** | **73.58(5.89)** | **8.24(0.76)** | **-0.99(0.49)** | **2.04(0.72)** |
| **Ion recognition-site** |  |  |  |  |  |  |
| **5 G:C 14 (W:W C)** | **35.09(23.0)** | **-43.67(83.39)** | **-8.57(41.78)** | **-4.45(4.81)** | **17.18(2.88)** | **3.56(2.17)** |
| **6 A:U 38 (W:W T)** | **-10.97(8.81)** | **11.29(9.66)** | **-24.32(9.27)** | **-4.25(0.97)** | **-1.53(0.87)** | **6.62(0.50)** |
| **7 U:** | **-77.23(9.22)** | **7.70(10.49)** | **42.95(10.20)** | **-1.30(0.85)** | **-3.18(0.78)** | **9.37(0.52)** |
| **8 G:C 47 (W:W C)** | **-1.52(2.78)** | **13.06(5.49)** | **32.81(3.82)** | **-0.30(0.47)** | **-1.93(0.32)** | **3.13(0.24)** |
| **40 A:U 48 (H:W T)** | **-56.43(59.3)** | **33.32(14.62)** | **15.52(16.38)** | **5.17(1.20)** | **-0.47(4.10)** | **8.53(2.74)** |
| **41 U:Cs:h T** | **26.85(6.49)** | **-27.76(19.75)** | **-25.51(7.65)** | **-3.20(1.29)** | **-3.46(0.49)** | **6.94(0.44)** |
| **42 G:C 13 (W:W C)** | **-2.71(2.47)** | **9.74(4.54)** | **28.67(3.27)** | **-0.23(0.37)** | **-1.91(0.31)** | **3.23(0.22)** |

**Table S3 (b): Mean and SD of Intra Base parameter (apo)**

| **Base pairing** | **Buckle** | **Open** | **Propeller** | **Stagger** | **Shear** | **Stretch** |
| --- | --- | --- | --- | --- | --- | --- |
| **Pseudoknot** |  |  |  |  |  |  |
| **2 G:C 17 (W:W C)** | **-2.64(7.50)** | **-0.01(3.08)** | **-4.62(6.91)** | **-0.14(0.29)** | **-0.17(0.25)** | **2.91(0.08)** |
| **3 G:C 16 (W:W C)** | **-10.02(8.20)** | **0.56(3.25)** | **-12.95(6.21)** | **-0.12(0.31)** | **0.08(0.31)** | **2.90(0.08)** |
| **4 C:G 15 (W:W C)** | **-9.62(8.09)** | **0.22(3.50)** | **-12.80(6.56)** | **0.08(0.32)** | **0.09(0.30)** | **2.91(0.09)** |
| **5 G:C 14 (W:W C)** | **-2.62(7.97)** | **-0.62(3.29)** | **-4.17(6.53)** | **0.003(0.31)** | **0.05(0.29)** | **2.93(0.09)** |
| **6 A:U 38 (W:W T)** | **-3.56(9.94)** | **-2.36(5.92)** | **-7.86(8.67)** | **-0.07(0.39)** | **-0.17(0.38)** | **2.91(0.15)** |
| **40 A:U 48 (H:W T)** | **-3.72(8.53)** | **0.83(5.71)** | **-10.42(6.34)** | **-0.02(0.40)** | **-0.16(0.27)** | **2.87(0.11)** |
| **Stem1** |  |  |  |  |  |  |
| **8 G:C 47 (W:W C)** | **-1.75(8.28)** | **-1.76(3.23)** | **-9.42(6.92)** | **-0.01(0.31)** | **-0.07(0.30)** | **2.92(0.09)** |
| **9 A:U 46 (W:W C)** | **-7.62(9.07)** | **2.55(6.11)** | **-13.01(8.88)** | **-0.24(0.40)** | **-0.21(1.07)** | **2.89(0.12)** |
| **10 G:C 45 (W:W C)** | **-9.06(8.36)** | **-0.57(3.49)** | **-8.62(7.18)** | **-0.23(0.31)** | **0.002(0.32)** | **2.91(0.09)** |
| **11 G:C 44 (W:W C)** | **-9.45(8.90)** | **1.26(3.67)** | **-10.55(6.76)** | **-0.19(0.30)** | **-0.08(0.30)** | **2.93(0.10)** |
| **12 C:G 43 (W:W C)** | **0.91(8.92)** | **1.34(4.46)** | **-9.32(6.74)** | **-0.05(0.32)** | **0.005(0.34)** | **2.94(0.11)** |
| **13 C:G 42 (W:W C)** | **0.98(7.60)** | **-0.41(3.33)** | **-10.25(6.44)** | **-0.14(0.30)** | **0.08(0.30)** | **2.91(0.09)** |
| **Stem2** |  |  |  |  |  |  |
| **24 G:C 37 (W:W C)** | **-0.90(8.68)** | **-0.88(3.34)** | **-5.85(6.92)** | **-0.01(0.31)** | **-0.11(0.31)** | **2.91(0.08)** |
| **25 C:G 36 (W:W C)** | **2.87(8.40)** | **0.26(3.48)** | **-8.41(6.57)** | **-0.01(0.31)** | **0.07(0.31)** | **2.91(0.09)** |
| **26 C:G 35 (W:W C)** | **2.04(7.97)** | **-0.12(3.49)** | **-8.85(6.70)** | **-0.16(0.29)** | **0.14(0.30)** | **2.91(0.09)** |
| **27 C:G 34 (W:W C)** | **1.38(8.11)** | **-0.34(3.46)** | **-6.73(7.18)** | **-0.14(0.31)** | **0.07(0.30)** | **2.92(0.09)** |
| **28 U:A 33 (W:W C)** | **9.24(9.37)** | **-1.09(5.37)** | **3.06(8.95)** | **0.06(0.41)** | **-0.07(0.28)** | **2.92(0.14)** |
| **Loop1** |  |  |  |  |  |  |
| **18 C:** | **0.0** | **0.0** | **0.0** | **0.0** | **0.0** | **0.0** |
| **19 A:** | **0.0** | **0.0** | **0.0** | **0.0** | **0.0** | **0.0** |
| **20 A:Cs:s C** | **31.22(8.30)** | **-36.38(6.23)** | **0.54(6.52)** | **-2.06(0.39)** | **1.09(0.28)** | **3.81(0.23)** |
| **21 A:G 3 (s:s T)** | **-29.67(7.66)** | **-61.69(4.57)** | **-26.05(6.16)** | **1.19(0.36)** | **-2.57(0.28)** | **2.42(0.20)** |
| **22 C:** | **0.0** | **0.0** | **0.0** | **0.0** | **0.0** | **0.0** |
| **23 U:** | **0.0** | **0.0** | **0.0** | **0.0** | **0.0** | **0.0** |
| **Loop2** |  |  |  |  |  |  |
| **29 G:** | **0.0** | **0.0** | **0.0** | **0.0** | **0.0** | **0.0** |
| **30 A:** | **0.0** | **0.0** | **0.0** | **0.0** | **0.0** | **0.0** |
| **31 A:** | **0.0** | **0.0** | **0.0** | **0.0** | **0.0** | **0.0** |
| **32 A:** | **0.0** | **0.0** | **0.0** | **0.0** | **0.0** | **0.0** |
| **Ion recognition-site** |  |  |  |  |  |  |
| **5 G:C 14 (W:W C)** | **-2.62(7.97)** | **-0.62(3.29)** | **-4.17(6.53)** | **0.003(0.31)** | **0.05(0.29)** | **2.93(0.09)** |
| **6 A:U 38 (W:W T)** | **-3.56(9.94)** | **-2.36(5.92)** | **-7.86(8.67)** | **-0.07(0.39)** | **-0.17(0.38)** | **2.91(0.15)** |
| **7 U:** | **0.0** | **0.0** | **0.0** | **0.0** | **0.0** | **0.0** |
| **8 G:C 47 (W:W C)** | **-1.75(8.28)** | **-1.76(3.23)** | **-9.42(6.92)** | **-0.01(0.31)** | **-0.07(0.30)** | **2.92(0.09)** |
| **40 A:U 48 (H:W T)** | **-3.72(8.53)** | **0.83(5.71)** | **-10.42(6.34)** | **-0.02(0.40)** | **-0.16(0.27)** | **2.87(0.11)** |
| **41 U:Cs:h T** | **11.65(29.60)** | **67.79(23.18)** | **59.27(20.43)** | **2.80(1.25)** | **2.69(1.35)** | **2.07(1.32)** |
| **42 G:C 13 (W:W C)** | **-0.98(7.60)** | **-0.41(3.33)** | **-10.25(6.44)** | **-0.14(0.30)** | **-0.08(0.30)** | **2.91(0.09)** |

**Table S3(c): The torsion and pseudo torsion angles for apo**

| **Base** | **CHI** | **PHASE** | **ETA** | **THETA** |
| --- | --- | --- | --- | --- |
| **Pseudoknot** |  |  |  |  |
| **2G** | **-156.72** \| **(57.13)** | **10.72** \| **(10.12)** | **159.31** \| **(42.04)** | **-147.95** \| **(21.48)** |
| **3G** | **-140.14** \| **(96.66)** | **7.71** \| **(9.95)** | **113.25** \| **(130.49)** | **-145.33** \| **(11.51)** |
| **4C** | **-160.75** \| **(30.89)** | **12.56** \| **(9.52)** | **119.35** \| **(124.53)** | **-146.78** \| **(7.13)** |
| **5G** | **-148.62** \| **(8.74)** | **18.58** \| **(11.73)** | **159.86** \| **(45.14)** | **30.30** \| **(9.00)** |
| **6A** | **-118.87** \| **(90.82)** | **11.83** \| **(16.50)** | **-105.40** \| **(7.75)** | **-150.75** \| **(30.59)** |
| **14C** | **-158.56** \| **(18.42)** | **21.78** \| **(11.32)** | **148.92** \| **(78.69)** | **-154.75** \| **(24.29)** |
| **15G** | **-158.98** \| **(32.10)** | **9.96** \| **(10.72)** | **159.34** \| **(44.81)** | **-144.05** \| **(9.69)** |
| **16C** | **-159.12** \| **(18.37)** | **12.59** \| **(9.51)** | **154.05** \| **(67.62)** | **-145.71** \| **(7.79)** |
| **17C** | **-160.01** \| **(22.12)** | **18.57** \| **(9.54)** | **150.57** \| **(76.13)** | **-157.71** \| **(13.53)** |
| **38U** | **-159.25** \| **(28.14)** | **15.71** \| **(10.40)** | **156.21** \| **(53.70)** | **-145.18** \| **(15.07)** |
| **40A** | **-110.18** \| **(13.67)** | **126.22** \| **(81.13)** | **-37.37** \| **(11.57)** | **-116.38** \| **(9.92)** |
| **48U** | **-159.71** \| **(21.58)** | **14.56** \| **(10.12)** | **161.30** \| **(20.87)** | **-147.15** \| **(12.08)** |
| **Stem1** |  |  |  |  |
| **8G** | **-99.06** \| **(139.99)** | **5.96** \| **(11.97)** | **-107.75** \| **(128.02)** | **-137.82** \| **(14.64)** |
| **9A** | **-158.44** \| **(30.63)** | **13.07** \| **(11.45)** | **151.47** \| **(64.93)** | **-146.58** \| **(14.61)** |
| **10G** | **-159.34** \| **(23.69)** | **16.05** \| **(10.91)** | **158.85** \| **(29.83)** | **-146.52** \| **(14.07)** |
| **11G** | **-159.36** \| **(28.09)** | **18.83** \| **(12.20)** | **148.01** \| **(79.11)** | **-147.02** \| **(14.52)** |
| **12C** | **-158.94** \| **(19.89)** | **19.10** \| **(11.23)** | **158.39** \| **(48.64)** | **-148.92** \| **(15.88)** |
| **13C** | **-159.05** \| **(17.13)** | **21.72** \| **(11.86)** | **154.56** \| **(65.90)** | **-152.75** \| **(16.41)** |
| **42G** | **87.74** \| **(148.88)** | **-45.10** \| **(14.27)** | **39.42** \| **(36.98)** | **-83.96** \| **(9.36)** |
| **43G** | **-158.26** \| **(29.30)** | **46.70** \| **(15.14)** | **137.33** \| **(9.10)** | **-158.57** \| **(28.28)** |
| **44C** | **-152.20** \| **(10.17)** | **25.79** \| **(12.83)** | **159.47** \| **(21.37)** | **-153.03** \| **(11.61)** |
| **45C** | **-155.00** \| **(10.18)** | **21.07** \| **(11.20)** | **160.16** \| **(33.18)** | **-145.52** \| **(10.60)** |
| **46U** | **-156.22** \| **(13.85)** | **20.12** \| **(12.06)** | **159.79** \| **(26.39)** | **-149.95** \| **(14.85)** |
| **47C** | **-158.93** \| **(21.80)** | **18.57** \| **(10.69)** | **155.39** \| **(60.78)** | **-146.36** \| **(9.18)** |
| **Stem2** |  |  |  |  |
| **24G** | **-52.593** \| **(164.68)** | **-9.66** \| **(16.48)** | **-128.09** \| **(92.46)** | **-126.12** \| **(15.79)** |
| **25C** | **-159.83** \| **(26.15)** | **14.76** \| **(12.04)** | **152.37** \| **(51.88)** | **-145.08** \| **(13.31)** |
| **26C** | **-159.66** \| **(21.69)** | **18.75** \| **(12.11)** | **155.61** \| **(59.80)** | **-148.44** \| **(14.67)** |
| **27C** | **-157.52** \| **(15.89)** | **22.77** \| **(13.37)** | **152.86** \| **(68.75)** | **-150.95** \| **(23.38)** |
| **28U** | **-151.69** \| **(11.17)** | **21.32** \| **(11.39)** | **159.47** \| **(41.70)** | **-138.89** \| **(10.03)** |
| **33A** | **-157.91** \| **(41.77)** | **9.12** \| **(9.49)** | **157.39** \| **(9.53)** | **-146.08** \| **(7.64)** |
| **34G** | **-159.51** \| **(34.01)** | **11.78** \| **(10.27)** | **153.33** \| **(69.91)** | **-149.63** \| **(10.24)** |
| **35G** | **-157.76** \| **(50.43)** | **11.61** \| **(10.16)** | **150.83** \| **(73.88)** | **-149.60** \| **(11.09)** |
| **36G** | **-155.29** \| **(60.67)** | **10.53** \| **(10.85)** | **138.61** \| **(97.84)** | **-143.86** \| **(9.55)** |
| **37C** | **-160.64** \| **(30.34)** | **14.38** \| **(10.46)** | **146.03** \| **(83.48)** | **-143.79** \| **(12.98)** |
| **Loop1** |  |  |  |  |
| **18C** | **-131.31** \| **(10.85)** | **138.93** \| **(67.87)** | **-154.19** \| **(7.49)** | **97.94** \| **(8.52)** |
| **19A** | **-141.1** \| **(68.93)** | **78.37** \| **(148.08)** | **-151.32** \| **(17.99)** | **147.50** \| **(22.92)** |
| **20A** | **-120.27** \| **(11.66)** | **30.57** \| **(44.52)** | **59.95** \| **(157.65)** | **101.30** \| **(129.27)** |
| **21A** | **-100.69** \| **(10.15)** | **-13.59** \| **(20.93)** | **-120.63** \| **(121.23)** | **-90.41** \| **(10.98)** |
| **22C** | **-129.19** \| **(28.14)** | **35.62** \| **(100.13)** | **155.60** \| **(39.16)** | **-78.84** \| **(131.61)** |
| **23U** | **-125.03** \| **(22.58)** | **37.92** \| **(24.24)** | **117.71** \| **(110.74)** | **86.31** \| **(47.06)** |
| **Loop2** |  |  |  |  |
| **29G** | **-149.54** \| **(11.10)** | **19.30** \| **(11.24)** | **157.37** \| **(15.79)** | **-125.34** \| **(14.48)** |
| **30A** | **-121.68** \| **(111.93)** | **1.59** \| **(20.11)** | **34.06** \| **(12.50)** | **-124.56** \| **(10.27)** |
| **31A** | **-153.03** \| **(35.47)** | **9.78** \| **(12.37)** | **139.84** \| **(77.63)** | **-128.60** \| **(76.42)** |
| **32A** | **-153.92** \| **(33.55)** | **9.73** \| **(10.08)** | **154.77** \| **(69.09)** | **-97.03** \| **(6.66)** |
| **Ion-Recognition-Site** |  |  |  |  |
| **5G** | **-148.62** \| **(8.74)** | **18.58** \| **(11.73)** | **159.86** \| **(45.14)** | **30.30** \| **(9.00)** |
| **6A** | **-118.87** \| **(90.82)** | **11.83** \| **(16.50)** | **-105.40** \| **(7.75)** | **-150.75** \| **(30.59)** |
| **7U** | **-128.78** \| **(12.47)** | **68.81** \| **(23.51)** | **-94.47** \| **(139.42)** | **-54.67** \| **(159.95)** |
| **8G** | **-99.06** \| **(139.99)** | **5.96** \| **(11.97)** | **-107.75** \| **(128.02)** | **-137.82** \| **(14.64)** |
| **40A** | **-110.18** \| **(13.67)** | **126.22** \| **(81.13)** | **-37.37** \| **(11.57)** | **-116.38** \| **(9.92)** |
| **41U** | **-121.85** \| **(42.28)** | **15.28** \| **(46.54)** | **139.42** \| **(87.29)** | **-106.38** \| **(59.29)** |
| **42G** | **87.74** \| **(148.88)** | **-45.10** \| **(14.27)** | **39.42** \| **(36.98)** | **-83.96** \| **(9.36)** |

**Table S4: Mean and SD of Overlap**

| **Base pairing** | **holo** |  | **apo** |  |
| --- | --- | --- | --- | --- |
|  | **Mean** | **SD** | **Mean** | **SD** |
| **Pseudoknot** |  |  |  |  |
| **2 G:C 17 (W:W C)** | **48.58** | **5.25** | **48.11** | **4.67** |
| **3 G:C 16 (W:W C)** | **48.14** | **4.72** | **44.58** | **6.92** |
| **4 C:G 15 (W:W C)** | **45.55** | **6.65** | **45.00** | **6.70** |
| **5 G:C 14 (W:W C)** | **45.55** | **6.65** | **48.11** | **4.67** |
| **6 A:U 38 (W:W T)** | **30.01** | **4.95** | **0.00** |  |
| **38 U:A 6 (W:W T)** | **30.01** | **4.95** | **0.00** |  |
| **40 A:U 48 (H:W T)** | **32.18** | **5.47** | **18.37** | **5.43** |
| **Stem1** |  |  |  |  |
| **8 G:C 47 (W:W C)** | **45.84** | **5.56** | **45.37** | **5.45** |
| **9 A:U 46 (W:W C)** | **43.73** | **6.14** | **47.22** | **5.46** |
| **10 G:C 45 (W:W C)** | **48.04** | **5.62** | **44.93** | **5.32** |
| **11 G:C 44 (W:W C)** | **47.66** | **4.94** | **49.13** | **5.16** |
| **12 C:G 43 (W:W C)** | **47.53** | **5.22** | **47.90** | **5.54** |
| **13 C:G 42 (W:W C)** | **47.28** | **5.64** | **44.58** | **6.92** |
| **Stem2** |  |  |  |  |
| **24 G:C 37 (W:W C)** | **47.80** | **4.80** | **49.02** | **5.21** |
| **25 C:G 36 (W:W C)** | **48.86** | **5.22** | **47.45** | **4.73** |
| **26 C:G 35 (W:W C)** | **47.37** | **5.66** | **42.57** | **6.19** |
| **27 C:G 34 (W:W C)** | **41.95** | **6.41** | **28.67** | **5.85** |
| **28 U:A 33 (W:W C)** | **27.61** | **6.07** | **0.00** |  |
| **Loop1** |  |  |  |  |
| **18 C:** | **0.00** |  | **0.00** |  |
| **19 A:** | **14.49** | **7.26** | **34.87** | **4.49** |
| **20 A:Cs:s C** | **17.94** | **3.29** | **5.78** | **4.30** |
| **21 A:G 3 (s:s T)** | **8.97** | **3.49** | **10.03** | **7.00** |
| **22 C:** | **12.13** | **5.03** | **0.00** |  |
| **23 U:** | **0.00** |  | **0.00** |  |
| **Loop2** |  |  |  |  |
| **29 G:** | **0.00** |  | **0.0** | **4.81** |
| **30 A:** | **21.06** | **4.83** | **21.02** | **4.02** |
| **31 A:** | **21.11** | **4.05** | **8.21** | **4.04** |
| **32 A:** | **9.89** | **5.14** | **42.57** | **6.19** |
| **Ion-Recognition-Site** |  |  |  |  |
| **5G** | **45.55** | **6.65** | **48.11** | **4.67** |
| **6A** | **30.01** | **4.95** | **0.00** |  |
| **7U** | **0.00** |  | **0.00** |  |
| **8G** | **49.33** | **5.52** | **31.48** | **5.00** |
| **40A** | **32.18** | **5.47** | **18.37** | **5.43** |
| **41U** | **0.00** |  | **0.0** |  |
| **42G** | **47.53** | **5.22** | **44.93** | **5.32** |

**Table S5 (a): Docking results from HDOCK**

| **Peptide-RNA complex** | **Docking Score** | **Confidence Score** | **Receptor interface residue pair** |
| --- | --- | --- | --- |
| ***Docked complex of GramidinD and Fluoride riboswitch*** | -393.89 | 0.99 | Stem2: 24,25,26,27,37  Loop1: 21,22,23,  Ion Recognition Site: 5,6,7, 41,42 |
| ***Docked complex of Magainin 2 and Fluoride riboswitch*** | -340.88 | 0.97 | Stem2: 24,25,26,27  Loop1: 18,19,20,21,22,23 |

**Table S5 (b): Docking results from HADDOCK**

| **Peptide-RNA complex** | **Cluster used in study** | **HADDOCK score** | **Cluster size** | **RMSD** | **Van der Waals energy** | **Electrostatic energy** | **Desolavation energy** | **Restraints violation energy** | **Buried Surface Area** | **Z-score** |
| --- | --- | --- | --- | --- | --- | --- | --- | --- | --- | --- |
| ***Chain A of Docked complex of Gramidin D and Fluoride riboswitch*** | **Cluster 4** | -90.9 +/- 10.0 | 4 | 11.5 +/- 0.7 | -11.8 +/- 1.3 | -376.9 +/- 39.6 | -6.2 +/- 7.3 | 24.1 +/- 1.5 | 1372.9 +/- 56.9 | -1.0 |
| ***Chain B of Docked complex of Gramidin D and Fluoride riboswitch*** | **Cluster 2** | -111.6 +/- 15. | 12 | 51.7 +/- 39.3 | -42.7 +/- 6.2 | -265.8 +/- 28.7 | -20.3 +/- 6.5 | 45.7 +/- 17.6 | 1521.4 +/- 173.3 | -1.3 |
| ***Docked complex of Magainin 2 and Fluoride riboswitch*** | **Cluster 1** | -110.8 +/- 4.5 | 21 | 9.4 +/- 0.3 | -44.9 +/- 3.4 | -410.5 +/- 23.2 | 3.1 +/- 6.6 | 31.0 +/- 3.9 | 1629.0 +/- 18.2 | -2.5 |
